# Genetic Analysis of Right Heart Structure and Function in 40,000 People

**DOI:** 10.1101/2021.02.05.429046

**Authors:** James P. Pirruccello, Paolo Di Achille, Victor Nauffal, Mahan Nekoui, Samuel N. Friedman, Marcus D. R. Klarqvist, Mark D. Chaffin, Shaan Khurshid, Carolina Roselli, Puneet Batra, Kenney Ng, Steven A. Lubitz, Jennifer E. Ho, Mark E. Lindsay, Anthony A. Philippakis, Patrick T. Ellinor

## Abstract

The heart evolved hundreds of millions of years ago. During mammalian evolution, the cardiovascular system developed with complete separation between pulmonary and systemic circulations incorporated into a single pump with chambers dedicated to each circulation. A lower pressure right heart chamber supplies deoxygenated blood to the lungs, while a high pressure left heart chamber supplies oxygenated blood to the rest of the body. Due to the complexity of morphogenic cardiac looping and septation required to form these two chambers, congenital heart diseases often involve maldevelopment of the evolutionarily recent right heart chamber. Additionally, some diseases predominantly affect structures of the right heart, including arrhythmogenic right ventricular cardiomyopathy (ARVC) and pulmonary hypertension. To gain insight into right heart structure and function, we fine-tuned deep learning models to recognize the right atrium, the right ventricle, and the pulmonary artery, and then used those models to measure right heart structures in over 40,000 individuals from the UK Biobank with magnetic resonance imaging. We found associations between these measurements and clinical disease including pulmonary hypertension and dilated cardiomyopathy. We then conducted genome-wide association studies, identifying 104 distinct loci associated with at least one right heart measurement. Several of these loci were found near genes previously linked with congenital heart disease, such as *NKX2-5, TBX3, WNT9B*, and *GATA4*. We also observed interesting commonalities and differences in association patterns at genetic loci linked with both right and left ventricular measurements. Finally, we found that a polygenic predictor of right ventricular end systolic volume was associated with incident dilated cardiomyopathy (HR 1.28 per standard deviation; P = 2.4E-10), and remained a significant predictor of disease even after accounting for a left ventricular polygenic score. Harnessing deep learning to perform large-scale cardiac phenotyping, our results yield insights into the genetic and clinical determinants of right heart structure and function.

---

The heart evolved hundreds of millions of years ago as a tubular organ^1^. Septation of the main pumping chamber of the heart into distinct left and right ventricles evolved later in birds, mammals, and some reptiles, and is under the control of conserved transcription factors such as *TBX5*^2^. Substantially greater delivery of oxygen to the systemic circulation—and to the heart itself—is the putative advantage of this separation of the circulatory system into a left heart-driven systemic circuit and a right heart-driven pulmonary circuit^3^.

The structures of the left and right heart are derived from different progenitor cell populations and operate under different pressure regimes: the left heart operates against high pressure, while the right heart generally faces little afterload. During embryogenesis, the left ventricle forms from the first heart field, while the right ventricle, the outflow tract, and portions of the atria form from the second heart field^4–7^. Septation of the outflow tract also requires neuroectodermal neural crest cells^8–10^.

The distinct embryological origins of the right and left ventricles likely explain, in part, the existence of right heart-predominant pathologies. These include arrhythmogenic right ventricular cardiomyopathy (ARVC)^11–14^, Brugada syndrome, and pulmonary hypertension. In addition, right ventricular dysfunction can play a role in other heart failure syndromes. The function of the right heart is an important determinant of outcomes in people who have heart failure with either reduced (HFrEF) or preserved left ventricular ejection fraction (HFpEF)^15–17^. HFpEF represents a heterogeneous set of diseases for which very few disease-modifying therapies exist. Consequently, there is substantial interest in identifying new therapies for conditions such as right ventricular dysfunction^18–21^.

The distinct pathologies, embryology, and physiology of the right heart motivated our efforts to quantify right heart structure and function, and to probe the common genetic basis for human variation in these measurements.

## Results

In this work, we developed deep learning models to determine the dimensions and function of the right atrium (RA), the right ventricle (RV), and the pulmonary artery (PA) in up to 45,000 UK Biobank participants. We then evaluated the epidemiologic associations, pathologic outcomes, and the common genetic basis of variation in these right heart structures.

### Reconstruction of right heart structures from cardiovascular magnetic resonance images

We first derived right heart measurements in the UK Biobank imaging substudy of over 45,000 people^22–24^ using deep learning models. To do so, a cardiologist created training data for deep learning models by manually tracing the right atrium and right ventricle in the four-chamber long axis view, and the right ventricle and pulmonary artery in the short axis view (**Figure 1**). This process, called semantic segmentation, yielded anatomical labels identifying the pixels belonging to cardiac structures in 714 short axis images and 445 four-chamber long axis images. Two U-Net derived deep learning models, containing long-range skip connections that allow for pixel-accurate segmentation, were then trained from these data: one for the four-chamber long axis view and another for the short axis views^25,26^. The deep learning models were then used to produce pixel labels for the remainder of the images. Quality assessment is detailed in the **Online Methods** and **Supplementary Note**.

**Figure 1:**
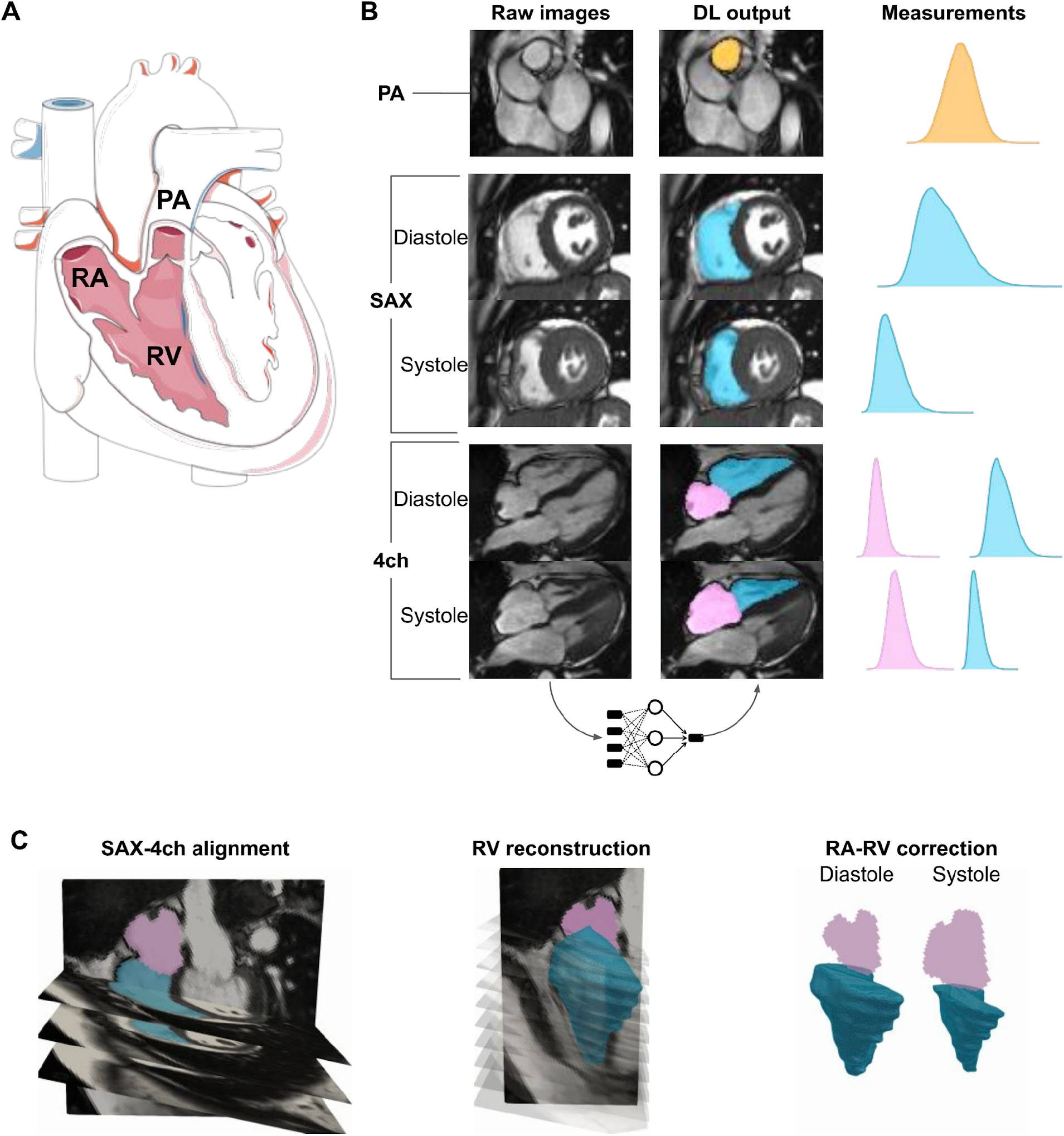
**Panel A**: Graphical depictions of the right heart structures in diastole and systole. RA = right atrium. PA = pulmonary artery. RV = right ventricle. In ventricular diastole, the tricuspid valve opens, allowing blood to flow from the right atrium into the right ventricle. The pulmonic valve is closed. In ventricular diastole, the right ventricle squeezes, closing the tricuspid valve and ejecting blood across the pulmonic valve into the pulmonary artery. During this time, the right atrium fills. The images in Panel A are derived from Servier Medical Art (licensed under creative commons by attribution). **Panel B**: Right heart structures and deep learning model output. PA = pulmonary artery. SAX = short axis view. 4ch = four-chamber long axis view. DL = deep learning. The pulmonary artery segmentation is colored in orange; the right ventricle is colored in blue; and the right atrium is colored in pink. The raw images on the leftmost panes are fed into the trained deep learning model, producing output that is colorized and laid on top of the raw images in the middle panes. This process is repeated for all participants and the output structures are measured, leading to population distributions of measurements as shown in the right panes. **Panel C**: Integration of SAX and 4ch data to reconstruct the right ventricle. The different images are aligned based on metadata provided from the MRI. A surface reconstruction technique is then applied (see **Methods** for details). Finally, reconstructed portions of the right ventricle that bulge into the right atrium are removed.

The deep learning model output was then post-processed to extract measurements of the right atrium, the right ventricle, and the pulmonary artery. The right atrium was only consistently visible in one view (the four-chamber long axis view), and therefore a 2-dimensional area was computed by summing the pixels and multiplying by their width and height. We computed the maximum and minimum area during the cardiac cycle, as well as the fractional area change (RA FAC), which is the ratio of the change in area between the maximum and minimum area divided by the maximum area.

The right ventricle has a complex 3-dimensional geometry; to estimate right ventricular structure, we integrated data from the short axis views and the four-chamber long axis view with a Poisson surface reconstruction approach, detailed in the **Online Methods**. We measured the maximum volume (right ventricular end diastolic volume; RVEDV), the minimum volume (right ventricular end systolic volume; RVESV), the difference between those two volumes (stroke volume), and the ejection fraction (RVEF).

The pulmonary trunk’s elliptical minor axis (diameter) was computed from short axis images at end-systole. For participants whose pulmonary trunk was visible in multiple short-axis slices, we refer to the component closest to the right ventricle as the pulmonary root, and the distal-most component as the proximal pulmonary artery.

In total, we were able to measure at least one right heart structure in 45,456 individuals, of whom 41,101 contributed to at least one genome-wide association study after genotyping quality control and exclusion for prevalent disease (**Table 1** and **Supplementary Figure 1**).

**Table 1:** Participant characteristics. displays characteristics of the 41,101 participants whose data contributed to at least one right heart phenotype GWAS. For quantitative phenotypes, values shown represent mean (SD). For count data, values shown represent count (%).

|  | Women | Men |
| --- | --- | --- |
| Participants | 21,932 | 19,169 |
| Age at time of MRI | 64 (8) | 65 (8) |
| BMI (kg/m<sup>2</sup>) | 26 (5) | 27 (4) |
| Height (cm) | 163 (6) | 176 (7) |
| Weight (kg) | 69 (13) | 83 (13) |
| Systolic Blood Pressure (mmHg) | 136 (19) | 142 (17) |
| Diastolic Blood Pressure (mmHg) | 77 (10) | 81 (10) |
| Drinking Status |  |  |
| Current | 20122 (92 %) | 18001 (94 %) |
| Never | 900 (4 %) | 439 (2 %) |
| Prefer not to answer | 7 (0 %) | 12 (0 %) |
| Previous | 746 (3 %) | 603 (3 %) |
| Standard drinks/week | 4.7 (5.3) | 5.6 (6.8) |
| Smoking status |  |  |
| Current | 605 (3 %) | 796 (4 %) |
| Never | 14334 (65 %) | 11289 (59 %) |
| Prefer not to answer | 85 (0 %) | 49 (0 %) |
| Previous | 6751 (31 %) | 6921 (36 %) |
| Smoking quantity (pack years) | 17 (14) | 21 (17) |
| Right atrium maximum area (cm<sup>2</sup>) | 22 (4) | 26 (5) |
| Right atrium minimum area (cm<sup>2</sup>) | 13 (3) | 16 (4) |
| Right atrium fractional area change (%) | 42 (6) | 40 (6) |
| Right ventricular end diastolic volume (mL) | 112 (21) | 155 (30) |
| Right ventricular end systolic volume (mL) | 44 (11) | 66 (17) |
| Right ventricular stroke volume (mL) | 68 (14) | 89 (19) |
| Right ventricular ejection fraction (%) | 61 (6) | 57 (6) |
| Proximal pulmonary artery diameter (cm) | 2.5 (0.3) | 2.7 (0.4) |
| Pulmonary artery root diameter (cm) | 2.5 (0.3) | 2.9 (0.3) |
| RA maximum area, BSA indexed (cm<sup>2</sup> / m<sup>2</sup>) | 13 (2) | 13 (2) |
| RA minimum area, BSA indexed (cm<sup>2</sup> / m<sup>2</sup>) | 7.3 (1.6) | 7.9 (1.8) |
| RV end diastolic volume, BSA indexed (mL / m<sup>2</sup>) | 64 (10) | 77 (14) |
| RV end systolic volume, BSA indexed (mL / m<sup>2</sup>) | 25 (6) | 33 (8) |
| RV stroke volume, BSA indexed (mL / m<sup>2</sup>) | 39 (7) | 44 (9) |
| Proximal PA diameter, BSA indexed (cm / m<sup>2</sup>) | 1.4 (0.2) | 1.3 (0.2) |
| PA root diameter, BSA indexed (cm / m<sup>2</sup>) | 1.4 (0.2) | 1.4 (0.2) |
| Left ventricular end diastolic volume (mL) | 123 (20) | 153 (28) |
| Left ventricular end systolic volume (mL) | 41 (11) | 58 (16) |
| Left ventricular stroke volume (mL) | 81 (12) | 95 (16) |
| Left ventricular ejection fraction (%) | 67 (5) | 63 (6) |

### Right heart structures are correlated with their left-heart counterparts

The mean and standard deviation of the right atrial area measurements, right ventricular volumes, and pulmonary artery diameters are described in **Table 1** and visualized in **Supplementary Figure 2**. Standard values aggregated by age bands and sex for each of the phenotypes are reported in **Supplementary Table 1**. The estimates of right atrial area from the four-chamber view are similar to those previously reported^27^, as are the proximal pulmonary artery diameters^28^. The estimates of right ventricular stroke volume are comparable to prior reports, but both end diastolic and end systolic volumes are approximately 10mL greater than those previously reported for steady-state free precession magnetic resonance imaging^29^. Consequently, the right ventricular ejection fraction estimates are proportionally lower than those of Foppa, *et al*.

We incorporated previously reported left ventricular traits and aortic traits^30,31^ in order to analyze cross-correlation between phenotypes of the right and left heart structures(**Supplementary Figure 3**). The volumetric measurements of the right and left ventricles were well correlated with one another (correlation between ventricular volumes was 0.84 at end-diastole and 0.71 at end-systole). In contrast, there was poorer correlation between right and left ventricular ejection fraction (correlation 0.48). This is consistent with drivers of contractility being only partially shared between the two ventricles, as well as multiplicative error due to the calculation of ejection fraction from two separately measured volumes. The ventricles nevertheless had well correlated stroke volumes (correlation 0.80), which is expected because stroke volume at steady-state is expected to be equal for both ventricles in the absence of valvular regurgitation or shunt.

The proximal pulmonary artery diameter was modestly correlated with right ventricular end systolic volume (correlation 0.49), suggesting shared right-heart related influences on the pulmonary artery diameter and right ventricular volumes. In addition, the pulmonary artery diameter and that of the ascending aorta—which share an embryological origin—were modestly correlated (correlation 0.45).

### Right heart measurements are associated with cardiovascular diseases

We tested PheCode-based disease definitions, which are derived from hospital diagnosis codes, for association with right heart phenotypes^32^. The right heart phenotypes were strongly correlated with atrial arrhythmias. The right atrial phenotypes were also associated with valvular diseases; the right ventricular phenotypes with obesity and heart failure; and the pulmonary artery phenotypes with obesity, blood pressure, and sleep disorders (**Figure 2, Supplementary Table 2**).

**Figure 2:**
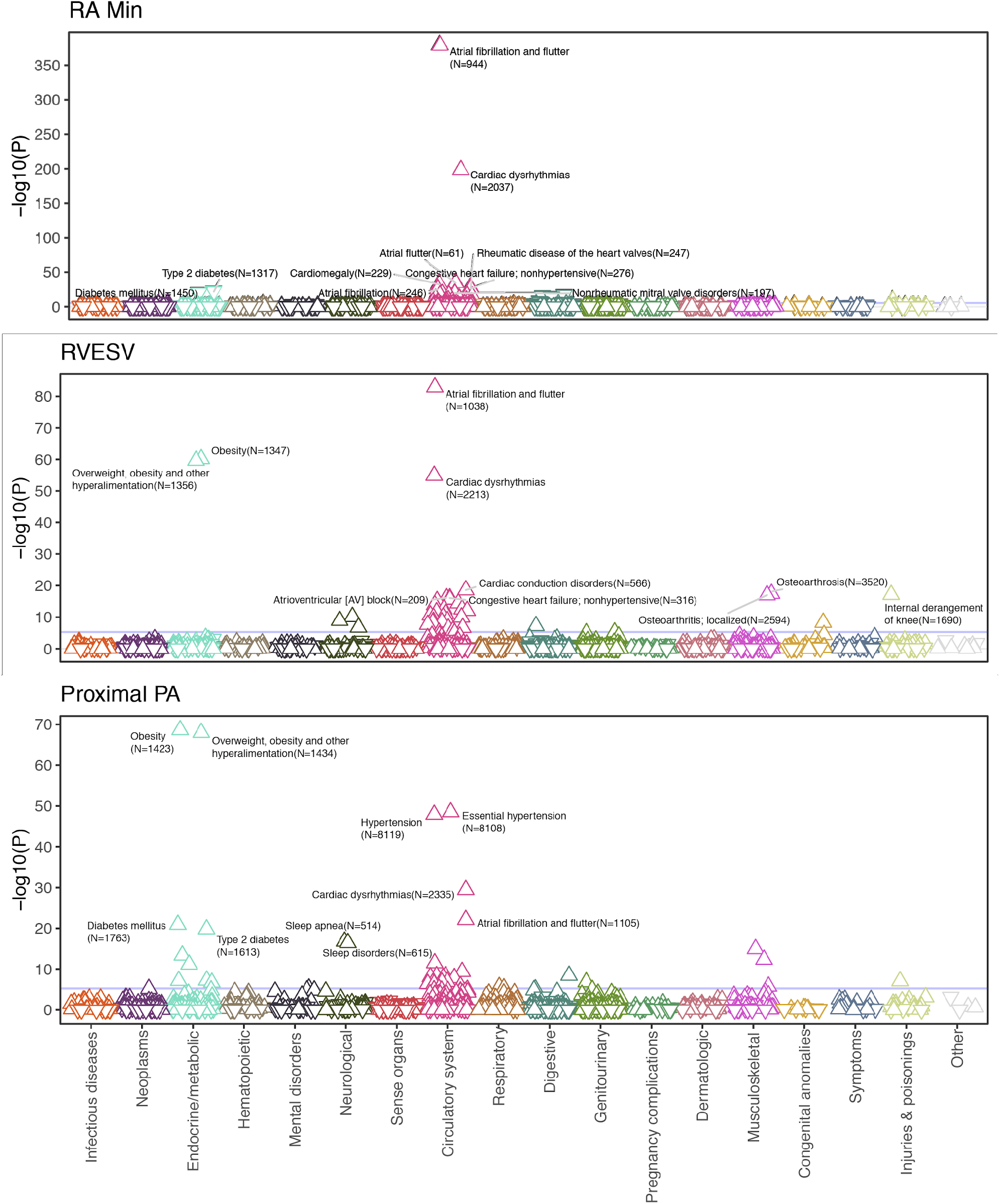
Right heart structures are associated with PheCode-based disease definitions. PheCode-based disease labels (**X-axis**) are plotted against a transformation of their association P value (**Y-axis**) with three right heart phenotypes: minimum right atrial area, right ventricular end systolic volume, and proximal pulmonary artery diameter. The modeled effect is a perturbation of the right heart trait among those with PheCode-based diseases identified prior to the time of magnetic resonance imaging, adjusting for anthropometric covariates and genetic principal components. The direction of the arrow indicates whether the presence of the disease is associated with an increase (upward arrow) or a decrease (downward arrow) of the right heart phenotype.

We also focused on three diseases with putative chamber-specific links to the right heart. We identified 1,033 individuals with a diagnosis of atrial fibrillation or flutter prior to undergoing MRI; 282 with congestive heart failure; and 21 with pulmonary hypertension (**Supplementary Table 3**). In a linear model, the right atrial FAC was 1.1 standard deviations (SD) lower among those with a history of atrial fibrillation or flutter than those without (P=2.6E-287). The RVEF was 0.51 SD lower among those with heart failure (P=6.6E-19). The proximal pulmonary artery diameter was 0.84 SD larger among those with pulmonary hypertension (P=5.9E-05). These findings confirmed expected structural correlations with prevalent cardiovascular diseases.

For two cardiovascular diseases—pulmonary hypertension and congestive heart failure—we modeled right ventricular volumes over the course of the cardiac cycle for individuals with and without disease (**Figure 3**). In these models, pulmonary hypertension (present in 21 participants) was associated with elevated volumes throughout the entire cardiac cycle, yielding a reduced RVEF. The excess volume that was attributable to disease accounted for as much as 34% of the total right ventricular volume (P = 1.8E-08) at end-systole and 15% (P = 1.4E-04) at end-diastole. Congestive heart failure (present in 282 participants) was also associated with elevated end-systolic volumes (14% elevation; P = 6.0E-17), but not with end-diastolic volumes (1% elevation; P = 0.44). As a negative control, 3,949 participants with cataract—a disease of the lens of the eye that is not expected to be linked to right ventricular size—was associated with no significant difference in right ventricular volumes compared to cataract-free individuals. These results demonstrate that different cardiovascular diseases can yield distinct perturbations of right ventricular volumes, and highlight the significant impact of pulmonary hypertension on right ventricular structure throughout the cardiac cycle in this population.

**Figure 3:**
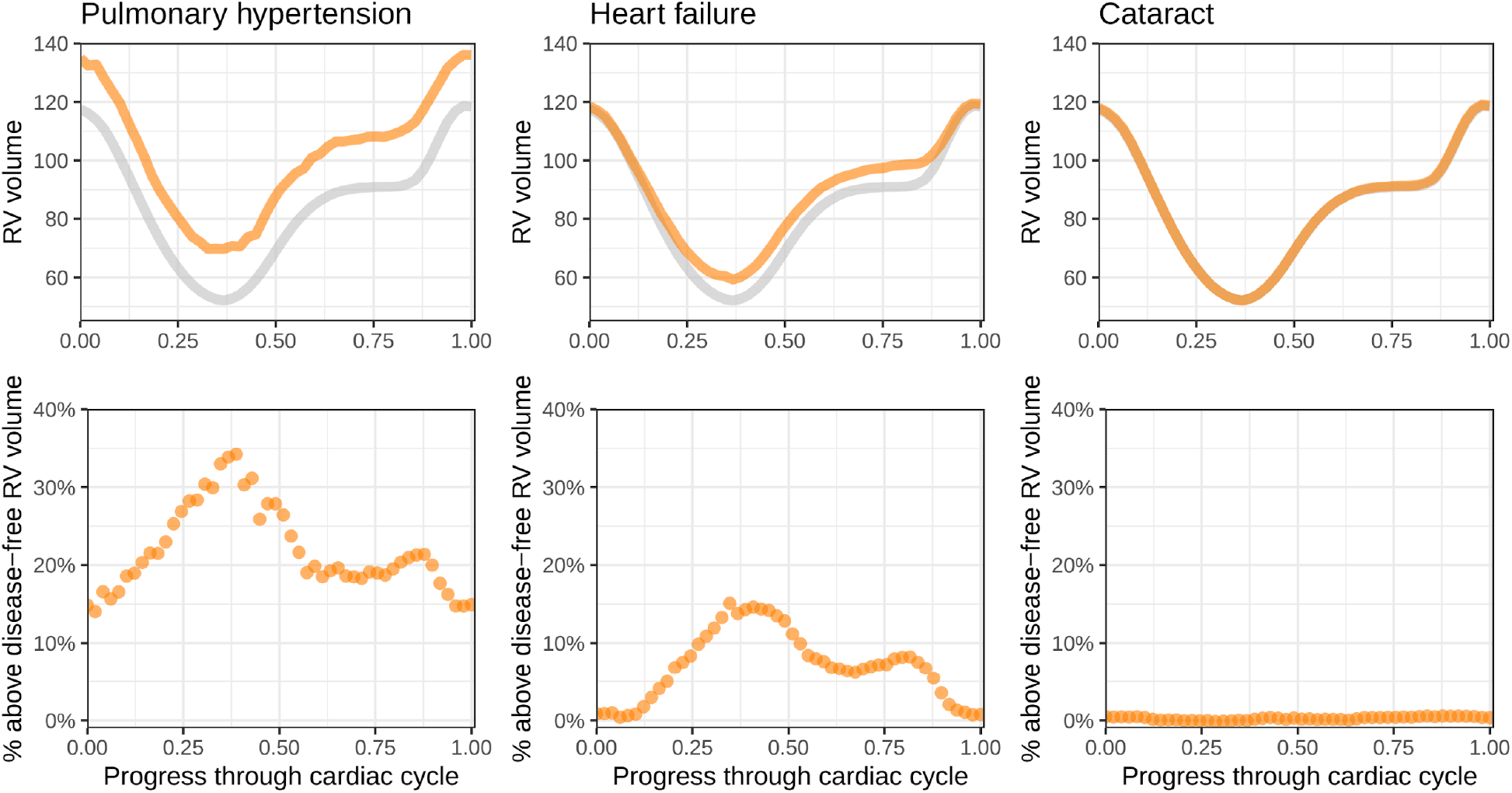
Perturbations to right ventricular volumes from prevalent diseases. **Top**: Disease diagnoses that occur prior to the date of MRI are linked with distinct changes in the volume of the right ventricle throughout the cardiac cycle. The x-axis represents fractions of a cardiac cycle (divided evenly into 50 components, starting at end-diastole). The y-axis represents volume in mL. Values are generated with a linear model for each time point; the gray line represents the population without disease, while the orange line represents the population with disease. In the UK Biobank, those with pulmonary hypertension have elevated RV volumes throughout the cardiac cycle, while those with heart failure predominantly have elevated end-systolic volumes. Cataract is used as a control to demonstrate little association between a non-cardiovascular disease and the volume of the right ventricle. **Bottom**: At each time, the right ventricular volume of individuals with disease is subtracted from the volume without disease and divided by the volume without disease. This represents the percentage above or below the disease-free right ventricular volume for those with disease.

### Right heart traits are heritable and genetically correlated with left heart traits

We then conducted genetic analyses of the right heart phenotypes. The size-related phenotypes showed significant heritability using BOLT-REML (as high as 0.37 for the maximum right atrial volume, 0.4 for right ventricular end-diastolic volume, and 0.42 for the pulmonary artery root diameter)^33,34^. Heritabilities were lower for measurements of right heart function, such as RVEF which had a heritability of 0.23.

We assessed genetic correlation between the right heart structures and previously reported left heart structures that include the left ventricle and the ascending aorta. Using individual-level data with BOLT-REML^34^, we found strong genetic correlation between the right and left ventricles (rg = 0.86 between RVEDV and LVEDV; rg = 0.75 between RVESV and LVESV; and rg = 0.61 between RVEF and LVEF). The proximal pulmonary artery diameter was most strongly correlated with the ascending aortic diameter (rg = 0.60). The genetic correlation matrix across all of the derived cardiovascular traits is available in **Supplementary Table 4** and **Supplementary Figure 4**, with trait heritabilities along the diagonal.

### Common genetic basis for the dimensions and function of the right heart

After establishing the heritability of the right heart traits, we conducted genome-wide association studies (GWAS) of each trait. We excluded participants with diagnoses of heart failure, atrial fibrillation, or myocardial infarction prior to their magnetic resonance imaging study (**Supplementary Figure 1**). We conducted nine primary GWAS: maximum and minimum right atrial area; RA FAC; RVESV, RVEDV, RVSV, and RVEF; pulmonary artery root diameter; and proximal pulmonary artery diameter. Up to 40,466 participants were included in these analyses, and we tested 11.6 million imputed SNPs with minor allele frequency (MAF) > 0.005 (**Table 2, Figure 4**). In addition, we evaluated the body surface area (BSA)-indexed versions of all traits except for RA FAC and RVEF (which are dimensionless), leading to a total of 16 GWAS (**Supplementary Table 5, Supplementary Figure 5**). Allowing loci to be counted once per trait, we identified 243 trait-locus pairs at a commonly used significance threshold of 5E-08. Accounting for multiple traits sharing loci, we identified 104 independent loci. Of these 104 loci, 66 were associated with at least two traits, and one locus (near *WNT9B*/*GOSR2*/*MYL4*) was associated with 12 right heart phenotypes. The greatest lambda GC was 1.20 from the BSA-indexed pulmonary artery root GWAS; *ldsc* revealed an intercept of 1.04, consistent with polygenicity rather than inflation (**Supplementary Table 6**)^35^. Six lead SNPs had Hardy-Weinberg equilibrium (HWE) P < 1E-06; re-analysis of those SNPs in a strictly European subset of samples resolved the HWE violations and yielded similar effect estimates (**Supplementary Table 7**).

**Table 2:**
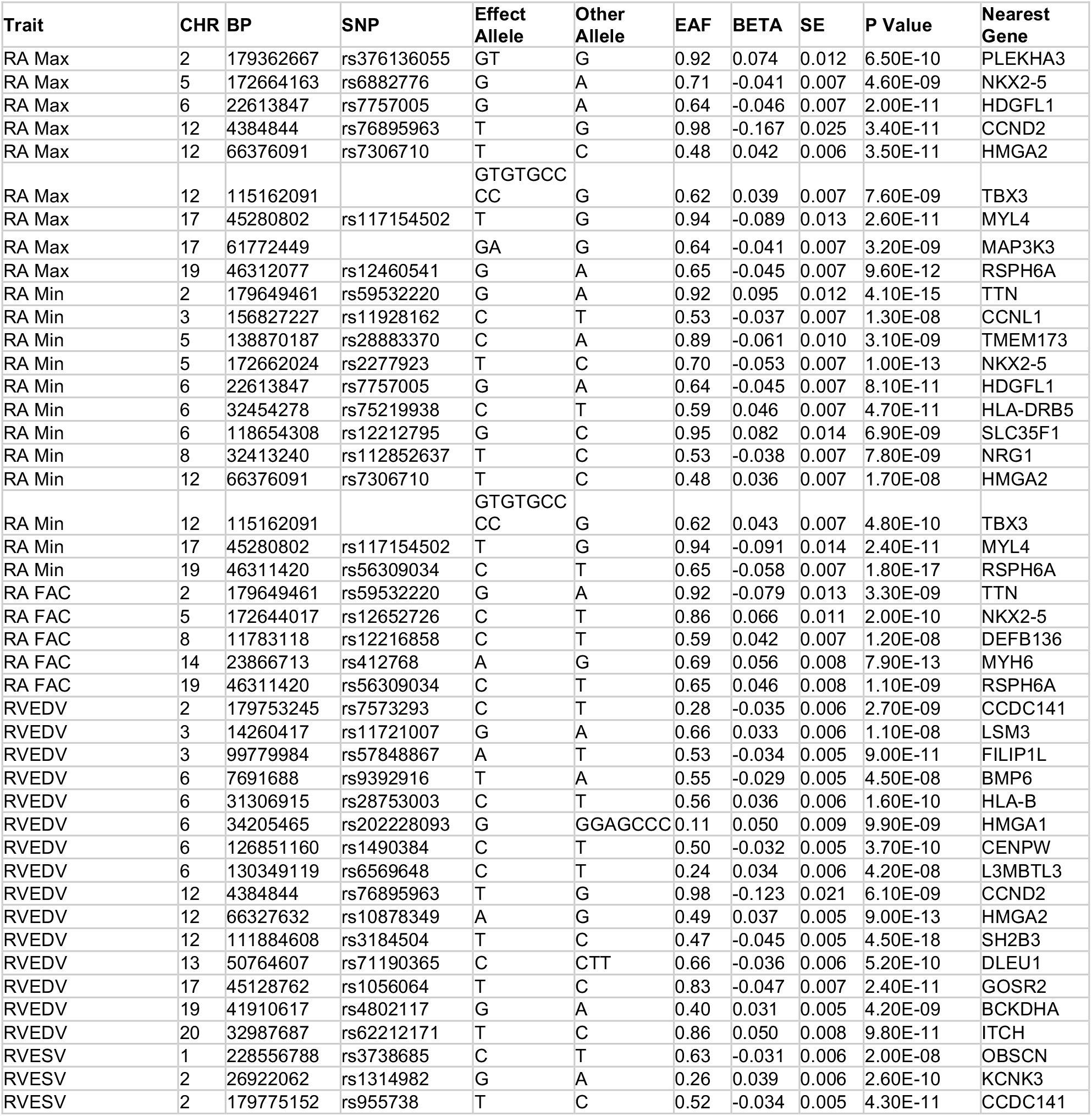
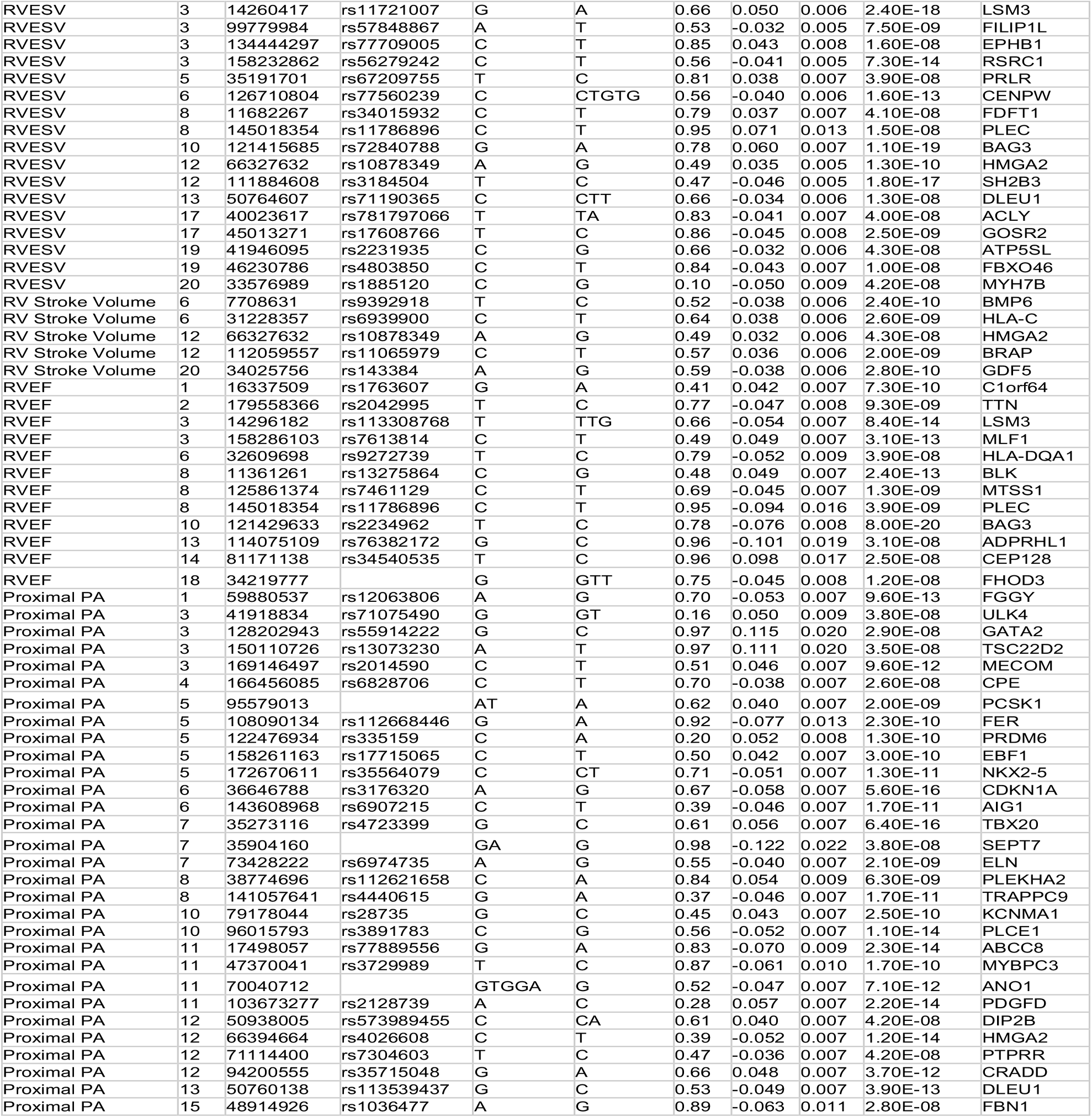
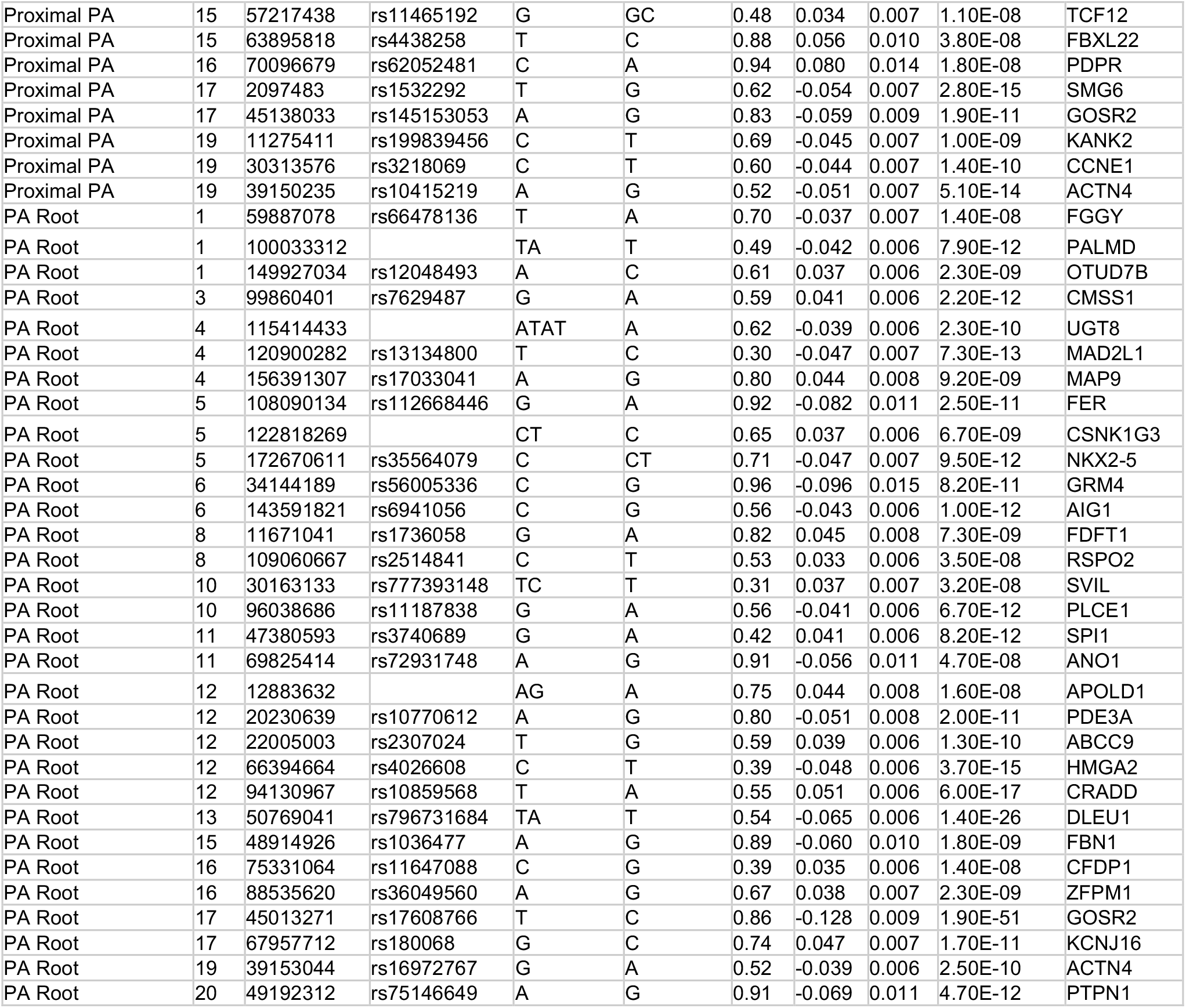
Lead SNPs. Lead SNPs from the right heart phenotypes. CHR: chromosome. BP: GRCh37 position. EAF: effect allele frequency. BETA: effect size. SE: standard error of effect size. Lead SNPs of the BSA-indexed phenotypes are listed in Supplementary Table 2.

**Figure 4:**
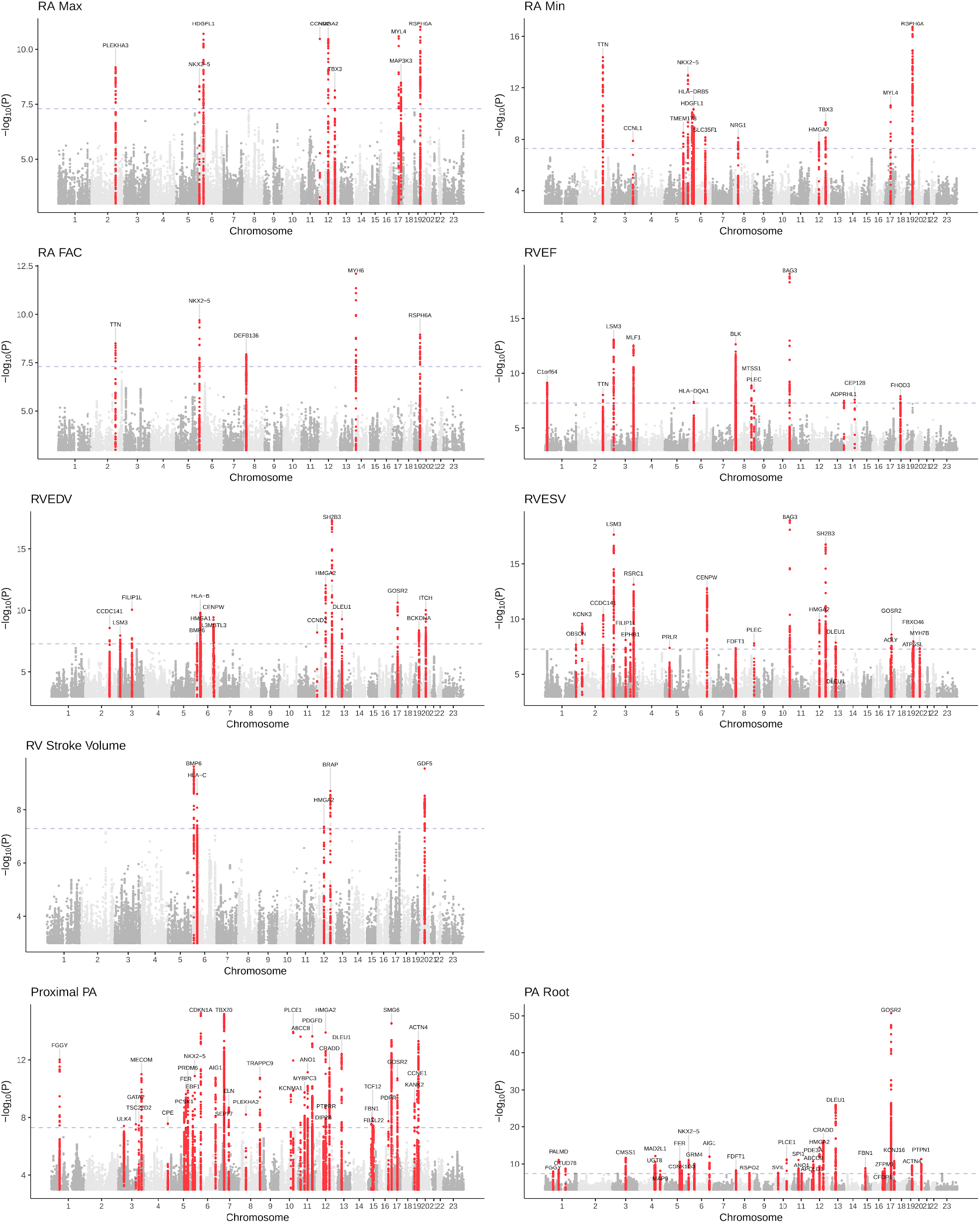
Manhattan plots. Manhattan plots show the chromosomal position (**X-axis**) and the strength of association (-log10 of the P value, **Y-axis**) for all non-BSA-indexed phenotypes. Loci that contain SNPs with P < 5E-08 are colored red and labeled with the name of the nearest gene.

To place the right heart results into context with prior work, we compared right heart loci with those previously associated with left ventricular and aortic dimensions (**Supplementary Figure 6**)^30,31^. Among the right ventricular phenotypes, the RVESV was linked with the greatest number of loci (20). Of these, seven loci had previously been associated with the RVESV’s left-heart counterpart (left ventricular end systolic volume; LVESV) at genome-wide significance. The *BAG3* locus is the most strongly associated with both RVESV and LVESV. Both traits shared the same lead SNP: rs72840788, which has a near perfect correlation with a SNP, rs2234962, that leads to the missense change p.Cys151Arg in the BAG3 protein (**Supplementary Figure 7**)^36^.

In contrast, at the *TTN* locus, the RVESV lead SNP (rs955738, GWAS P = 4.3E-11) was in linkage equilibrium (r^2^ = 0.001) with the LVESV lead SNP (rs2562845, GWAS P = 1.3E-23). However, both SNPs were among the secondary signals for these traits: for LVESV, rs955738 was associated with P = 1.9E-11; for RVESV, rs2562845 was associated with P = 4.2E-08 (**Supplementary Figure 8**). It is possible that this distinction between primary association signals in the two ventricles is associated with differences in the regulation of *TTN* between the first (LV) and second (RV) heart fields, but establishing this will require additional investigation.

Among loci that were significant only for RVESV and not for LVESV, some, like the *GATA4*/*CTSB* locus, had a cluster of sub-threshold SNPs for LVESV. At this locus, the strongest LVESV-associated SNP (rs7012446, P = 1.5E-06) was weakly correlated (r^2^ = 0.16) with the RVESV lead SNP (rs34015932, P = 4.1E-08), also suggesting allelic heterogeneity (**Supplementary Figure 9**). Other loci, such as that of *OBSCN* (encoding obscurin, a giant sarcomeric protein in the same family as titin), appeared to be right-ventricle specific, showing very little evidence of association with the left ventricle (**Supplementary Figure 10**).

### TWAS highlights role of WNT signaling in pulmonary root diameter

Across all phenotypes, the strongest GWAS association was between the pulmonary root diameter and rs17608766 (P = 1.9E-51), near *GOSR2*. In a transcriptome-wide association study (TWAS) based on gene expression data from the aorta from GTEx v7^37^, at the *GOSR2* locus we observed an association between pulmonary root diameter and *WNT9B* (full results in **Supplementary Table 8**). Interestingly, *WNT9B* is expressed in the endocardium overlying the heart valves during development, and loss of *WNT9B* leads to defective valve formation^38^. This locus was also recently shown to be linked with the mitral valve annular diameter^39^.

The strongest TWAS association for the proximal pulmonary artery diameter was with *PDGFD*, which is also the nearest gene to the lead SNP rs2128739. *PDGFD* loss-of-function variants were recently implicated in pulmonary hypertension in a sequencing-based case-control study^40^.

### Chamber-specific cell type enrichment

To identify relevant cell types most relevant for the right atrial and right ventricular phenotypes, we performed stratified linkage disequilibrium (LD) score regression analysis that integrated single nucleus RNA-sequencing data from Tucker *et al*^41^. The strongest enrichment was seen between RVEF and right ventricular cardiomyocytes, while the strongest enrichment for the right atrial phenotypes was for vascular smooth muscle cell-like nuclei (**Supplementary Figure 11**).

### Rare variant association test

Up to 13,523 individuals with imaging data had exome sequencing performed in the first batch of 50,000 exomes in the UK Biobank. After accounting for multiple testing, loss of function variants in one gene (*AAGAB*) had significant association with the proximal pulmonary artery diameter (diameter larger by 0.42cm on average among the 14 individuals with *AAGAB* loss-of-function variants; P = 1.2E-06; **Supplementary Figure 12**). The AAGAB protein is involved in clathrin-mediated endocytosis^42^, and haploinsufficiency of *AAGAB* has previously been reported to be associated with punctate palmoplantar keratoderma^43,44^. These prior reports are noteworthy in this context because of the association between palmoplantar keratoderma and ARVC^45,46^. Nevertheless, we did not identify an association between *AAGAB* variants and RVEF (P=0.06) or RVESV (P=0.28). Additionally, common variants at the *AAGAB* locus demonstrated no significant association with pulmonary artery diameter or right ventricular size and function in the GWAS. Future studies in additional populations will be required to assess the significance of the observed association between *AAGAB* and right heart phenotypes.

### GWAS loci enriched in uncommon and difficult to phenotype cardiac diseases

To investigate the association between loci identified in this study and diseases that are not well represented (such as congenital heart diseases) or difficult to identify due to lack of specific diagnostic codes in the electronic health record (such as arrhythmogenic right ventricular cardiomyopathy), we performed proximity-based testing to assess enrichment of gene sets near the GWAS loci. We identified disease-related gene sets using the Open Targets platform (gene lists in **Supplementary Table 9**)^47^ and asked whether more of those genes than expected by chance were found within 500kb of the GWAS lead SNPs. Note that because the number of permutations generated by SNPSnap in the following tests was 10,000, the strongest possible association P value was 1.0E-04^48^.

The right atrial loci were in proximity to six Open Targets atrial septal defect-related genes (*ACE, DMPK, MIR208A, NKX2-5, PLN, TBX5*) with one-tailed permutation P = 1.0E-04. The right ventricular GWAS loci were in proximity to six ARVC-related genes (*DSP, JUP, PPP1R13L, RBM20, TMEM43, TTN*) with P = 1.1E-03. And the pulmonary artery loci were in proximity to 15 conotruncal abnormality-linked genes (*BAZ1B, CEP152, DYNC2H1, ELN, EPHB4, FBN1, GATA4, KCNJ8, MECOM, NKX2-5, PDE3A, PDE5A, PLCE1, RYR1, SMARCA4*) with P = 3.0E-04 (**Supplementary Figure 13**).

We also analyzed a previously described panel of 129 cardiomyopathy-linked genes to contrast RVESV loci with LVESV loci^30^. Five of these genes were within a 500kb radius of the RVESV loci; of these, three (*BAG3, TMEM43*, and *TTN*) had previously been found near genome-wide significant LVESV loci, while two (*GATA4* and *JUP*) were not (**Supplementary Figures 9** and **14**). The RVESV and LVESV associations at the *GATA4* locus have been described above. *JUP*, the gene that encodes plakoglobin, is a desmosomal protein that has also been associated with arrhythmogenic right ventricular cardiomyopathy and palmoplantar keratoderma, a syndrome known as Naxos disease^11,13,49^.

### Right heart polygenic scores are linked with cardiomyopathy and atrial fibrillation

Finally, we assessed the association between polygenic scores derived from the right heart GWAS and incident cardiovascular diseases in UK Biobank participants unrelated to the individuals who underwent MRI.

A polygenic score for RVESV was associated with dilated cardiomyopathy (680 events and 409,944 non-events; HR 1.28 per SD; P = 2.4E-10; **Figure 5**). Notably, even after adjustment for the previously reported left ventricular end systolic volume BSA-indexed polygenic score^30^, the RVESV polygenic score remained associated with DCM (HR 1.17 per SD, P = 6.9E-05). Because of imprecision in clinical phenotyping from electronic health records (EHR), future work will be required to understand whether the RVESV polygenic score identifies additional cases of DCM that are driven by right ventricular dysfunction, or whether the score is identifying shared drivers of right- and left-ventricular dysfunction that were not ascertained in the left ventricular GWAS.

**Figure 5:**
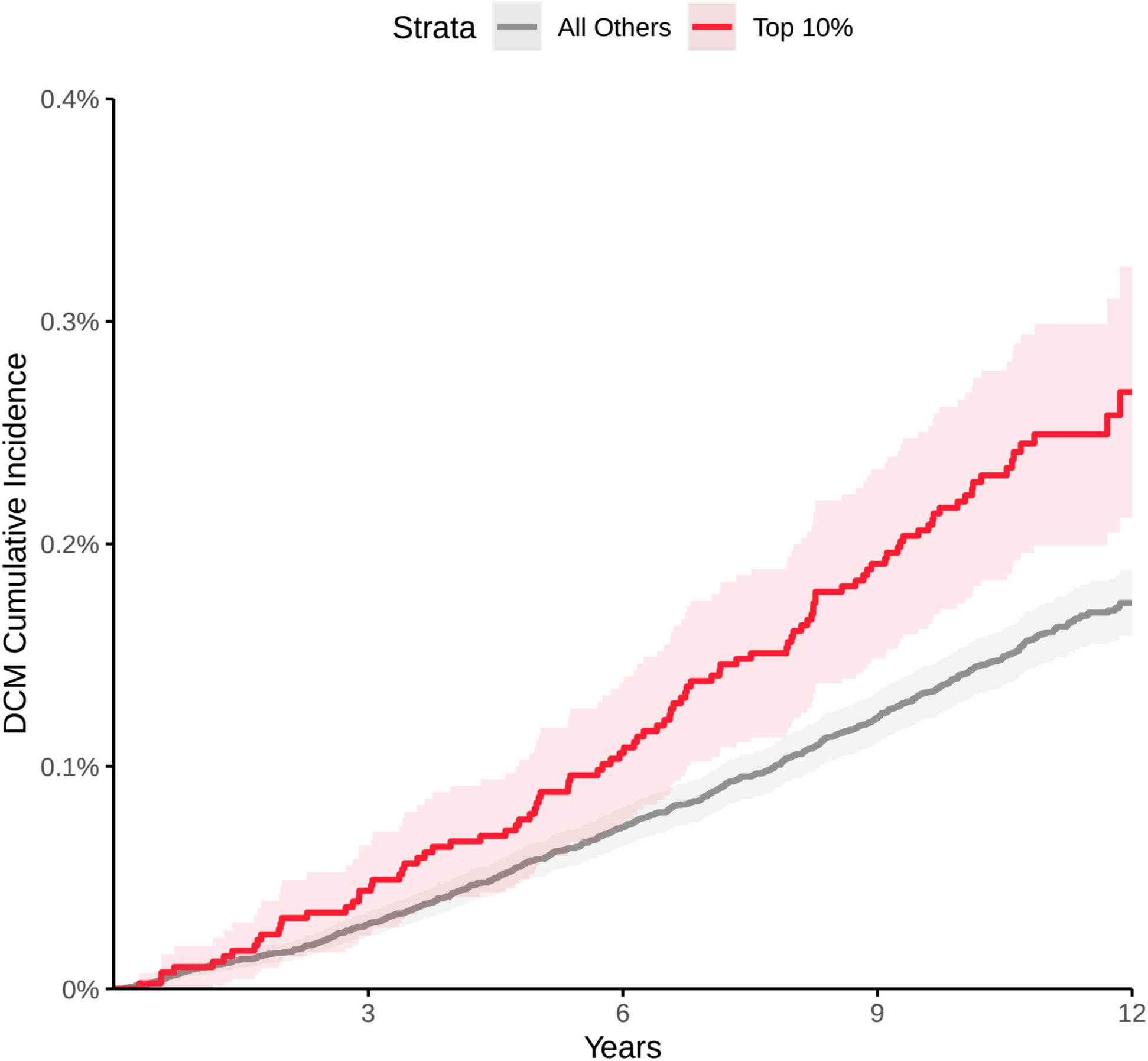
Cumulative incidence of dilated cardiomyopathy stratified by genetic prediction of RVESV. Individuals unrelated within 3 degrees of the participants who underwent MRI in the top 10% for the RVESV PRS (red) and bottom 90% (gray). **X-axis**: years since enrollment in the UK Biobank. **Y-axis**: cumulative incidence of dilated cardiomyopathy. Those in the top 10% of genetically predicted RVESV had an increased risk of DCM (Cox HR 1.53, P = 8E-05) compared with those in the bottom 90% in up to 12 years of follow-up time after UK Biobank enrollment.

A polygenic score for the fractional area change of the right atrium in the four-chamber view was weakly inversely associated with the risk of atrial fibrillation or flutter (for 15,122 events and 402,951 non-events; HR 0.98 per SD; P = 1.9E-03). Results were similar when considering only atrial flutter as the outcome of interest (927 atrial flutter events and 423,824 non-events; HR 0.90 per SD; P = 1.2E-03). We did not find a link between the risk of pulmonary hypertension (1,582 incident events and 423,136 non-events) and genetic predictions of proximal pulmonary artery diameter or pulmonary root diameter (P=0.59 and 0.93, respectively).

### Limitations

This study is subject to several limitations. All data were derived from deep learning models of short axis or four-chamber long axis views from cardiovascular magnetic resonance imaging. These models have imprecision that would be reduced with further training data. Like any deep learning model, these models can fail and produce non physiologic measurements when presented with images that contain features not seen in the training data. An advantage of the semantic segmentation approach in this work is that outliers can be visually inspected and the model re-trained as needed. The right atrial measurements are two-dimensional estimates of a three-dimensional structure and therefore cannot capture complete information about atrial volume. The short axis images have a coarse 10mm slice thickness which leads to partial volume imaging, which can be particularly difficult to visualize at the apex of the right ventricle, leading to under- or over-estimation by the deep learning model. Although we have attempted to correct for this by incorporating the higher resolution four-chamber long axis data during the surface reconstruction process, the correction itself can introduce additional artifacts: the image acquisitions for the short axis measurements are not simultaneous with one another or with the four-chamber long axis measurements, which can create misalignment (e.g., due to differences in breath holding) that introduces error when reconstructing the right ventricle. The deep learning models have not been tested outside of the specific devices and imaging protocols used by the UK Biobank and may not generalize to other data sets without additional fine-tuning. Participants’ cardiac rhythm at the time of MRI (particularly normal sinus rhythm versus atrial fibrillation) was not adjudicated. The study population is largely of European ancestry, similar to the remainder of UK Biobank, limiting generalizability of the findings to other populations. The individuals who underwent MRI in the UK Biobank tend to be healthier than the remainder of the UK Biobank population, which itself is healthier than a general population. Finally, because we have used hospital-based ICD codes and procedural codes to identify individuals with disease, our study lacks an ARVC-specific analysis, and our disease definitions are susceptible to misclassification.

## Discussion

We produced measurements of the right heart, including the right atrium, right ventricle, and pulmonary artery; analyzed their relationships with one another and with cardiovascular diseases; and identified 104 distinct genetic loci that are associated with these right-heart measurements. We drew several conclusions from these findings.

First, right heart phenotypes, including structural and functional measurements of the right atrium, right ventricle, and pulmonary artery, are heritable. While they share strong epidemiological and genetic correlation with the corresponding left heart structures, our findings of partial genetic correlation and distinct genome-wide significant loci also imply distinct drivers of variation between right and left heart structures. Developing a better understanding of these distinct drivers may ultimately permit more targeted therapies for right ventricle-predominant heart failure syndromes and primary cardiomyopathies such as ARVC.

Second, we found that the GWAS loci were enriched for genes associated with developmental diseases such as atrial septal defect and conotruncal defects. In addition to the GWAS loci addressed above, several others were notable for connections to cardiovascular development. Right heart structures were associated with SNPs near *NKX2-5*, which plays a key role in maintaining the progenitor pool of cells of the secondary heart field^5^; *TBX3*, which controls the formation of the sinus node and loss of which leads to outflow tract malformations and septal defects^50,51^; and *MYL4*, which encodes atrial light chain 1, missense variants in which have been linked to familial atrial fibrillation^52^.

Third, we observed links between right ventricular measurements—and polygenic predictions of these measurements—and disease. Individuals with pre-existing diagnoses of heart failure had reduced RVESV at the time for MRI, and those with pulmonary hypertension had markedly enlarged right ventricular volumes throughout the cardiac cycle (**Figure 3**). In the remainder of the population that was unrelated to those who underwent MRI, a polygenic predictor of RVESV was a strong predictor of a diagnosis of dilated cardiomyopathy (**Figure 5**). Notably, the RVESV polygenic score remained a significant predictor of dilated cardiomyopathy even after accounting for a previously reported genetic prediction of the left ventricle—implying a genetic basis for the role of right ventricular dysfunction in the pathogenesis of dilated cardiomyopathy. Consistent with emerging clinical evidence, this suggests that right ventricular structure and function are not merely of anthropomorphic interest, but actually represent endophenotypes for cardiomyopathy.

Fourth, despite our observation of strong epidemiological association between pulmonary hypertension and the proximal pulmonary diameter, we did not find an association between the polygenic predictor of proximal pulmonary diameter and the incidence of pulmonary hypertension. This lack of association may be because pulmonary hypertension in the UK Biobank is largely environmentally driven; may indicate that the genetic contributions to pulmonary artery diameter in disease-free individuals are not significantly associated with pulmonary artery pressure; or may be due to weak instrument bias.

Finally, machine learning enables the derivation of complex traits in a manner that is scalable. This permits biobank-scale investigation of previously understudied human phenotypes, such as measurements of the right atrium, right ventricle, and pulmonary artery; and promises to accelerate our understanding of cardiovascular disease.

## Online Methods

### Study design

Except where otherwise stated, all analyses were conducted in the UK Biobank, which is a richly phenotyped, prospective, population-based cohort that recruited 500,000 individuals aged 40-69 in the UK via mailer from 2006-2010^24^. We analyzed 487,283 participants with genetic data who had not withdrawn consent as of February 2020. Access was provided under application #7089 and approved by the Partners HealthCare institutional review board (protocol 2019P003144).

Here we provide an overview of the methods used in this manuscript that are explained in more detail below. We manually annotated pixels from magnetic resonance images from the UK Biobank: the pulmonary artery and the left and right ventricles were annotated in the short axis view, and the right atrium and right ventricle were annotated in the four-chamber long axis view. We then trained two deep learning models (one for each of the views) with our manual annotations, and applied this model to the remaining images in the UK Biobank. For the right ventricle, we integrated the data from the four chamber view and the short axis view to generate a surface mesh and derived the ventricular volumes from this mesh. We analyzed the relationships between each of these derived quantitative measurements of the right heart. We also analyzed their relationships with diseases and other phenotypes in the UK Biobank. Then, we excluded people with known disease and conducted genome-wide association studies of the right heart phenotypes. We performed transcriptome-wide association studies (TWAS) that incorporated publicly available gene expression data with our GWAS results to prioritize genes at most genomic loci. We analyzed the GWAS results in light of the four-chamber single nucleus sequencing data that is publicly available. We also performed a rare-variant association test in up to 13,523 UK Biobank participants with both imaging and exome sequencing data. Polygenic scores produced from SNPs associated with right heart phenotypes in the UK Biobank GWAS were used to predict incident atrial fibrillation or flutter, dilated cardiomyopathy, and pulmonary hypertension in the UK Biobank participants whose data did not contribute to the GWAS.

Statistical analyses were conducted with R version 3.6 (R Foundation for Statistical Computing, Vienna, Austria).

### Cardiovascular magnetic resonance imaging protocols

At the time of this study, the UK Biobank had released images in over 45,000 participants of an imaging substudy that is ongoing^22,23^. Cardiovascular magnetic resonance imaging was performed with 1.5 Tesla scanners (Syngo MR D13 with MAGNETOM Aera scanners; Siemens Healthcare, Erlangen, Germany), and electrocardiographic gating for synchronization^23^. Several cardiac views were obtained. For this study, two views (the long axis four-chamber view and the short axis view) were used. In both of these views, balanced steady-state free precession cines, consisting of a series of 50 images throughout the cardiac cycle for each view, were acquired for each participant^23^. For the four-chamber images, only one imaging plane was available for each participant, with an imaging plane thickness of 6mm and an average pixel width and height of 1.83mm. For the short axis view, several imaging planes were acquired. Starting at the base of the heart, 8mm-thick imaging planes were acquired with approximately 2mm gaps between each plane, forming a stack perpendicular to the longitudinal axis of the left ventricle to capture the ventricular volume. For the short axis images, the average pixel width and height was 1.86mm.

### Semantic segmentation and deep learning model training

Semantic segmentation is the process of assigning labels to pixels of an image. Here, we labeled pixels within specific anatomical structures (the right atrial blood pool, the right ventricular blood pool, and the pulmonary artery blood pool), using a process similar to that described in our prior work evaluating the thoracic aorta^31^. Segmentation of cardiovascular structures was manually annotated in four-chamber and short axis images from the UK Biobank by a cardiologist (JP). To produce the model used in this manuscript, 714 short axis images were chosen, manually segmented, and used to train a deep learning model with PyTorch and fastai v1.0.61^25,53^. The same was done separately with 445 four-chamber images. For both views, the models were based on a U-Net-derived architecture constructed with a ResNet34 encoder that was pre-trained on ImageNet^26,54–56^. The Adam optimizer was used^57^. The models were trained with a cyclic learning rate training policy^58^. 80% of the samples were used to train the model, and 20% were used for validation. Held-out test sets that were not used for training or validation were used to assess the final quality of both models.

Two separate models were trained: one for the short axis images, and one for the four-chamber long axis images. The hyperparameters used in their training are described below. For both models, random perturbations of the input images (“augmentations”) were applied, including affine rotation, zooming, and modification of the brightness and contrast.

For the short axis images, all images were resized initially to 104×104 pixels during the first half of training, and then to 224×224 pixels during the second half of training. The model was trained with a mini-batch size of 16 (with small images) or 8 (with large images). Maximum weight decay was 1E-03. The maximum learning rate was 1E-03, chosen based on the learning rate finder^25,59^. Because the right ventricle and pulmonary artery blood pools occupied very little of the overall short axis image area, a focal loss function was used (with alpha 0.7 and gamma 0.7), which can improve performance in the case of imbalanced labels^60^. When training with small images, 60% of iterations were permitted to have an increasing learning rate during each epoch, and training was performed over 30 epochs while keeping the weights for all but the final layer frozen. Then, all layers were unfrozen, the learning rate was decreased to 1E-07, and the model was trained for an additional 10 epochs. When training with large images, 30% of iterations were permitted to have an increasing learning rate, and training was done for 30 epochs while keeping all but the final layer frozen. Finally, all layers were unfrozen, the learning rate was decreased to 1E-07, and the model was trained for an additional 10 epochs.

For the four-chamber long axis images, all images were resized initially to 76×104 pixels during the first half of training, and then to 150×208 pixels during the second half of training. The model was trained with a mini-batch size of 4 (with small images) or 2 (with large images). Maximum weight decay was 1E-02. Cross entropy loss was used^61^. 30% of iterations were permitted to have an increasing learning rate during each epoch. When training with small images, the maximum learning rate was initially 1E-03, and training was performed over 50 epochs while keeping all weights frozen except for the final layer. Then, all layers were unfrozen, the learning rate was decreased to 3E-05, and the model was trained for an additional 15 epochs. When training with large images, the maximum learning rate was set to 3E-04, and the model was trained for 50 epochs while keeping all but the final layer frozen. Finally, all layers were unfrozen, the learning rate was decreased to 1E-07, and the model was retrained for an additional 15 epochs.

The final short axis and four-chamber long axis models were then applied, respectively, to all available short axis images and four-chamber long axis images available in the UK Biobank as of November 2020.

### Deep learning model output quality control

Accuracy of the two deep learning models was assessed with additional manually annotated images that were not used for model training or validation, with each annotation category (such as pulmonary artery blood pool) evaluated based on the Sørensen-Dice coefficient^62,63^, which scales from 0 (no agreement between manual and automated annotations) to 1 (perfect agreement). Images with no pixels assigned to a feature by either the truth labels or the deep learning model output were assigned to have a Dice coefficient of 1.

### Right atrial measurements from the four-chamber long axis view

Three long-axis views were obtained in the UK Biobank (two-chamber, three-chamber, and four-chamber). Of these, only the four-chamber view reliably captures the right atrium. We therefore treated the right atrium as a planar surface, counting the pixels that were labeled by the four chamber semantic segmentation model as right atrium, and multiplying that number by the height and width of each pixel to obtain a right atrial area (with units of cm^2^). For each individual, we obtained the maximum atrial area, the minimum atrial area, and the fractional area change (maximum area minus minimum area, divided by maximum area).

### Pulmonary artery measurements from the short axis view

For most individuals in the UK Biobank, the pulmonary artery can be readily visualized in the basal-most short axis imaging planes. When the pulmonary artery was visible in multiple imaging planes, we measured the artery in both the basal-most and the apical-most plane that still captured a pulmonary artery cross-section. To facilitate reproducibility, we only evaluated images from the frame representing ventricular end-systole. We refer to the basal-most segment of pulmonary artery as the “proximal pulmonary artery” and the apical-most segment that sits just basal to the right ventricular outflow tract as the “pulmonary root.”

The proximal pulmonary artery and pulmonary root were treated as ellipses. We computed major and minor axes using classical image moment algorithms^31,64^. For both the proximal pulmonary artery and the pulmonary root, the length of the minor elliptical axis (i.e., the diameter) was computed. We excluded any measurements where the artery was divided into more than one connected component^65^. For the proximal pulmonary artery, we permitted elliptical eccentricity values of up to 0.86 (where eccentricity is 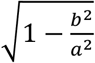, with *a* being the elliptical major axis and *b* being the elliptical minor axis). We permitted a liberal eccentricity cutoff because a common cause of high eccentricity in the proximal pulmonary artery images was out-of-plane curvature of the proximal pulmonary artery, which erroneously elongates the major elliptical axis but does not significantly affect the minor elliptical axis (which we are using as the diameter). For the pulmonary artery root, we required an eccentricity below 0.77. We used a more stringent cutoff at the root because a common cause of high eccentricity in these images was partial imaging of the right ventricle, which can erroneously foreshorten the minor elliptical axis. In addition, we excluded images where the cross-sectional area of the pulmonary artery was less than 2 cm^2^.

### Right ventricular annotation and surface reconstruction integrating long- and short-axis data

The right ventricle was visible in multiple views, permitting a reconstruction of its volume through the incorporation of data from multiple images. Stacked together, images from the short axis view provided a complete 3-dimensional representation of the right ventricle; however, this stack had a coarse 10mm resolution along the length of the right ventricle from base to apex. In contrast, the four-chamber long axis view had approximately 2mm resolution along the same axis. To take advantage of the strengths of both sources of data, we reconstructed the surface of the right ventricle using a Poisson surface reconstruction technique described in detail below. This enabled the computation of right ventricular end diastolic volume, end systolic volume, stroke volume, and ejection fraction.

To produce consistent estimates of the RV volumes throughout the cardiac cycle, we integrated information from the long- and short-axis segmentations by reconstructing 3-dimensional surfaces enclosing the RV cavity. We first used image metadata from the standard *Image Position (Patient) [0020,0032]* and *Image Orientation (Patient) [0020,0037]* DICOM tags to co-rotate the 4-chamber and short-axis slices into the same reference system. Then, we implemented a custom reconstruction routine based on the Poisson algorithm to generate surfaces that fitted through the boundaries of the RV segmentations^66^. As the Poisson algorithm requires local curvature as an input, we specified for the surface normal directions to lie onto the plane of the MRI slices and to be locally oriented towards either the pericardium (at the free wall) or the left ventricle (at the interventricular septum). The reconstructed RVs were then post-processed to correct for eventual artifacts in the basal short-axis slices, where the segmentation model may occasionally mistake the right atrium for part of the RV. Leveraging the fine resolution of the long-axis CMR in the apex-to-base direction, we discarded the portions of the reconstructed RVs that overextended above the plane separating the long-axis segmentations of the right atrium and of the RV (i.e., approximately co-aligned with the tricuspid valve plane). Finally, the RV volumes were estimated from the reconstructed surfaces using a discrete version of the divergence theorem, as implemented in the open-source VTK library (Kitware Inc.).

### Phenotypic characterization of right heart structure and function

Using R version 3.6, we evaluated the mean and standard deviation of the right heart measurements, described them in age- and sex-stratified tables, and created sex-stratified kernel density plots with *ggplot2*^67^. We computed the Pearson correlation between all right heart phenotypes and available left heart phenotypes that were previously described^30,31^.

We tested for association between the right heart phenotypes and PheCode-based disease labels derived from ICD-10 codes and OPCS-4 codes that were present prior to each participant’s magnetic resonance imaging date^32^. Association between each disease code and the right heart phenotypes was performed with linear models accounting for the MRI serial number, sex, the first five principal components of ancestry, age at enrollment, the cubic natural spline of age at the time of MRI, and the genotyping array. Splines were not placed on age at enrollment because of its collinearity with age at the time of MRI.

We used three custom disease definitions to focus on chamber-specific disease relationships (atrial fibrillation with RA FAC; heart failure with RVESV; and pulmonary hypertension with the proximal pulmonary artery diameter; defined in **Supplementary Table 3**). Association between each disease (as a binary independent variable) and the right heart phenotypes (as the dependent variable) was performed using a linear model that also accounted for the MRI serial number, sex, the first five principal components of ancestry, age at enrollment, the cubic natural spline of age at the time of MRI, and the genotyping array. As above, splines were not placed on age at enrollment because of its collinearity with age at the time of MRI.

We also modeled the association between three diseases (pulmonary hypertension, heart failure, and cataract) and right ventricular volume throughout the cardiac cycle. The magnetic resonance images were acquired as a series of 50 images throughout a cardiac cycle, and so our Poisson surface reconstruction yielded right ventricular volume for each of these timepoints (one-fiftieth of a cardiac cycle). At each of these timepoints, we used a linear model to test the association between the right ventricular volume (independent variable) and the presence or absence of each of the three diseases, as well as covariates that included the heart rate at the time of MRI, weight, height, age at enrollment, the cubic natural spline of age at the time of MRI, sex, genotyping array, and the first five principal components of ancestry. In the results, we report the P value for the linear model regression coefficient for the disease. To model the estimated volume for individuals with or without disease, we compute the output of the linear model for a 55-year-old woman who enrolled 5 years previously in the UK Biobank, 162 centimeters tall and weighing 75.6 kilograms. We then toggle the presence or absence of disease in the model to obtain volumes with or without disease, fixing other covariates.

### Genotyping, imputation, and genetic quality control

UK Biobank samples were genotyped on either the UK BiLEVE or UK Biobank Axiom arrays and imputed into the Haplotype Reference Consortium panel and the UK10K+1000 Genomes panel^68^. Variant positions were keyed to the GRCh37 human genome reference. Genotyped variants with genotyping call rate < 0.95 and imputed variants with INFO score < 0.3 or minor allele frequency <= 0.005 in the analyzed samples were excluded. After variant-level quality control, 11,631,796 imputed variants remained for analysis.

Participants without imputed genetic data, or with a genotyping call rate < 0.98, mismatch between self-reported sex and sex chromosome count, sex chromosome aneuploidy, excessive third-degree relatives, or outliers for heterozygosity were excluded from genetic analysis^68^. Participants were also excluded from genetic analysis if they had a history of pulmonary hypertension, atrial fibrillation, heart failure, or coronary artery disease documented by ICD code or procedural code from the inpatient setting prior to the time they underwent cardiovascular magnetic resonance imaging at a UK Biobank assessment center. Our definitions of these diseases in the UK Biobank are provided in **Supplementary Table 3**.

### Heritability and genome-wide association analyses

We analyzed nine primary right heart phenotypes. For the right atrium, we assessed maximum area, minimum area, and fractional area change. For the right ventricle, we assessed end diastolic volume, end systolic volume, stroke volume, and ejection fraction. For the pulmonary system, we assessed the diameter of the pulmonary root and the proximal pulmonary artery. In addition, we analyzed body surface area-indexed values for all areas and volumes (i.e., excluding RA FAC and RVEF which are dimensionless). In total, we conducted 16 genome-wide association studies with these traits.

BOLT-REML v2.3.4 was used to assess the SNP-heritability of the phenotypes, as well as their genetic correlation with one another using the directly genotyped variants in the UK Biobank^33^.

Before conducting genome-wide association studies, a rank-based inverse normal transformation was applied to the quantitative right heart traits^69^. All traits were adjusted for age at enrollment, age and age^2^ at the time of MRI, the first 10 principal components of ancestry, sex, the genotyping array, and the MRI scanner’s unique identifier.

Genome-wide association studies for each phenotype were conducted using BOLT-LMM version 2.3.4 to account for cryptic population structure and sample relatedness^33,34^. We used the full autosomal panel of 714,558 directly genotyped SNPs that passed quality control to construct the genetic relationship matrix (GRM), with covariate adjustment as noted above. Associations on the X chromosome were also analyzed, using all autosomal SNPs and X chromosomal SNPs to construct the GRM (N=732,193 SNPs), with the same covariate adjustments and significance threshold as in the autosomal analysis. In this analysis mode, BOLT treats individuals with one X chromosome as having an allelic dosage of 0/2 and those with two X chromosomes as having an allelic dosage of 0/1/2. Variants with association P < 5·10^−8^, a commonly used threshold, were considered to be genome-wide significant. In addition, we used a secondary threshold of P < 3.1·10^−9^ (5·10^−8^ divided by 16 phenotypes) to identify associations that were study-wide significant.

We identified lead SNPs for each trait. Linkage disequilibrium (LD) clumping was performed with PLINK-1.9^70^ using the same participants used for the GWAS, rather than a generic reference panel. We outlined a 5-megabase window (--clump-kb 5000) and used a stringent LD threshold (--r2 0.001) in order to account for long LD blocks such as those near the Williams-Beuren locus on chromosome 7 and the Noonan syndrome locus on chromosome 12^71–73^. With the independently significant clumped SNPs, distinct genomic loci were then defined by starting with the SNP with the strongest P value, excluding other SNPs within 500kb, and iterating until no SNPs remained. Independently significant SNPs that defined each genomic locus are termed the lead SNPs.

Lead SNPs were tested for deviation from Hardy-Weinberg equilibrium (HWE) at a threshold of P < 1E-06^70^. To assess whether the HWE violations affected the association signals, SNPs with HWE P < 1E-06 were re-analyzed with R’s *glm* after excluding samples that were not within the UK Biobank’s centrally-adjudicated “white British” subset, using the same covariates as the BOLT-LMM model.

Linkage disequilibrium (LD) score regression analysis was performed using *ldsc* version 1.0.0^35^. With *ldsc*, the genomic control factor (lambda GC) was partitioned into components reflecting polygenicity and inflation, using the software’s defaults.

### Transcriptome-wide association study

For each phenotype, we performed a TWAS to identify correlated genes based on imputed cis-regulated gene expression^74–76^. We used FUSION with eQTL data from GTEx v7. Precomputed transcript expression reference weights for the aorta (used for the pulmonary artery traits), left ventricle (used for the right ventricular traits), and right atrial appendage (used for the right atrial traits) were obtained from the FUSION authors’ website (http://gusevlab.org/projects/fusion/)^37,75^. FUSION was then run with its default settings.

### Stratified LD Score Regression

To identify putative cell types most relevant for each GWAS trait, we performed stratified linkage disequilibrium (LD) score regression analysis using single nucleus RNA-sequencing data from Tucker *et al*^35,41,77^. Cell type specific markers within the RA and RV were calculated separately for the 9 main cell types using a *limma-voom* differential expression model on aggregated counts per individual^78^. Only individuals with greater than 25 nuclei of a given cell type were considered. Genes were sorted by t statistic per cell type and the top 90% of genes were used to generate LD Score Regression annotations^77^. SNPs within 100 KB of any gene from a specific cell type were annotated for the respective cell type using 1000 Genomes European individuals^79^. We then performed stratified LD score regression with these annotations in combination with the baseline model described in Finucane, *et al*, 2015, only including high quality HapMap3 SNPs^80^. We used the RA cell type specific annotations and RV cell type specific annotations for the RA and RV specific GWAS traits, respectively.

### Exome sequencing

We conducted an exome sequencing analysis in the first 50,000 exomes released by the UK Biobank. Exome sequencing was performed by Regeneron and reprocessed centrally by the UK Biobank following the Functional Equivalent pipeline^81^. Exomes were captured with the IDT xGen Exome Research Panel v1.0, and sequencing was performed with 75-base paired-end reads on the Illumina NovaSeq 6000 platform using S2 flowcells. Alignment to GRCh38 was performed centrally with BWA-mem^82^. Variant calling was performed centrally with GATK 3.0^83^. Variants were hard-filtered if the inbreeding coefficient was < -0.03, or if none of the following were true: read depth was greater than or equal to 10; genotype quality was greater than or equal to 20; or allele balance was greater than or equal to 0.2. Variants were annotated with the Ensembl Variant Effect Predictor version 95 using the --pick-allele flag^84^. LOFTEE 1.0 was used to identify high-confidence loss of function variants: stop-gain, splice-site disrupting, and frameshift variants^85^. In total, 49,997 exomes were available, of which a subset overlapped with the participants who had undergone magnetic resonance imaging: 12,420 with right atrial measurements, 12,699 with right ventricular measurements, and 13,523 pulmonary artery measurements.

### Rare variant association test

We conducted a collapsing burden test to assess the impact of loss-of-function variants in up to 13,523 participants who had undergone exome sequencing data and magnetic resonance imaging. Variants with MAF >= 0.001 were excluded. We excluded genes with fewer than 10 loss-of-function variants passing the above criteria. The models testing for association between loss-of-function in each gene and the right heart traits were adjusted for weight (kg), height (cm), body mass index (kg/m^2^), the MRI serial number, age at enrollment, the cubic natural spline of age at the time of MRI, sex, the genotyping array, and the first five principal components of ancestry. As above, splines were not placed on age at enrollment because of its collinearity with age at the time of MRI.

### Open Targets gene set enrichment at GWAS loci

Using the Open Targets platform, we created gene sets corresponding to ARVC, atrial fibrillation, atrial septal defect, and transposition of the great arteries (TGA) or conotruncal anomaly by fetching all genes with an overall association score of 0.05 or greater (**Supplementary Table 9**)^47^. Using SNPsnap, we generated 10,000 sets of SNPs that matched the lead SNPs based on parameters including minor allele frequency, SNPs in linkage disequilibrium, distance from the nearest gene, and gene density. We counted the number of OpenTargets genes within 500kb of the lead SNPs from our study. We then repeated the same procedure for each of the 10,000 synthetic SNPsnap lead SNP lists, to set a neutral expectation for the number of overlapping genes based on chance. This allowed us to compute one-tailed permutation P values for each group of disease genes (with the most extreme possible P value based on 10,000 randomly chosen sets of SNPs being 1·10^−4^).

### Polygenic risk analysis

We computed a polygenic score based on 21 clumped, genome-wide significant SNPs (of which 20 were lead SNPs) from the RVESV GWAS. We applied this score to the entire UK Biobank population, after excluding any participant who had undergone imaging or who was related within 3 degrees to individuals with imaging. We analyzed the relationship between this polygenic prediction of the RVESV and dilated cardiomyopathy using a Cox proportional hazards model as implemented by the R *survival* package^86^. We excluded individuals with disease that was diagnosed prior to enrollment in the UK Biobank. We counted survival as the number of years between enrollment and disease diagnosis (for those with disease) and those being censored due to death, loss to follow-up, or end of follow-up time. We adjusted for covariates including sex, the cubic basis spline of age at enrollment, the interaction between the cubic basis spline of age at enrollment and sex, the genotyping array, the first five principal components of ancestry, and the cubic basis splines of height (cm), weight (kg), BMI (kg/m^2^), diastolic blood pressure, systolic blood pressure.

The same procedure was performed to produce a 5-SNP polygenic score for the RA fractional area change (all of which were lead SNPs) that was tested for association with atrial fibrillation and flutter. And the procedure was repeated to produce a 42-SNP polygenic score for the pulmonary artery diameter (of which 38 were lead SNPs) that was tested for association with pulmonary hypertension. The SNP weights are available in **Supplementary Table 10**.

To assess the impact of the RVESV PRS even after accounting for the previously reported LVESVi PRS, we also added the LVESVi PRS as a covariate to the RVESV-dilated cardiomyopathy Cox model.

## Data availability

UK Biobank data are made available to researchers from research institutions with genuine research inquiries, following IRB and UK Biobank approval. GWAS summary statistics will be available upon publication at the Broad Institute Cardiovascular Disease Knowledge Portal (http://www.broadcvdi.org). All other data are contained within the article and its supplementary information, or are available upon reasonable request to the corresponding author.

## Code availability

The code used to perform Poisson surface reconstruction from segmentation output is located at https://github.com/broadinstitute/ml4h and is available under an open-source BSD license. The code used to perform permutation testing to assess enrichment of disease-related genes near GWAS loci is located at https://github.com/carbocation/genomisc and is available under an open-source BSD license.

## Author contributions

JPP and PTE conceived of the study. JPP and VN annotated images. JPP trained the deep learning models. PD performed surface reconstruction. JPP, VN, MDC, PD, SNF, and MDRK conducted bioinformatic analyses. JPP and PTE wrote the paper. All other authors contributed to the analysis plan or provided critical revisions.

## Sources of funding

This work was supported by the Fondation Leducq (14CVD01), and by grants from the National Institutes of Health to Dr. Ellinor (1RO1HL092577, R01HL128914, K24HL105780) and Dr. Ho (R01HL134893, R01HL140224, K24HL153669). This work was supported by a John S LaDue Memorial Fellowship and the Sarnoff Cardiovascular Research Foundation Scholar Award to Dr. Pirruccello. Dr. Nauffal is supported by NIH grant 5T32HL007604-35. Dr. Khurshid is supported by NIH grant T32HL007208. Dr. Lubitz is supported by NIH grant 1R01HL139731 and American Heart Association 18SFRN34250007. This work was supported by a grant from the American Heart Association Strategically Focused Research Networks to Dr. Ellinor. Dr. Lindsay is supported by the Fredman Fellowship for Aortic Disease and the Toomey Fund for Aortic Dissection Research. This work was funded by a collaboration between the Broad Institute and IBM Research.

## Disclosures

Drs. Pirruccello has served as a consultant for Maze Therapeutics. Dr. Batra is supported by grants from Bayer AG and IBM applying machine learning in cardiovascular disease. Dr. Lubitz receives sponsored research support from Bristol Myers Squibb / Pfizer, Bayer AG, Boehringer Ingelheim, and Fitbit, and has consulted for Bristol Myers Squibb / Pfizer and Bayer AG, and participates in a research collaboration with IBM. Dr. Ng is employed by IBM Research. Dr. Ho is supported by a grant from Bayer AG focused on machine-learning and cardiovascular disease and a research grant from Gilead Sciences. Dr. Ho has received research supplies from EcoNugenics. Dr. Philippakis is employed as a Venture Partner at GV; he is also supported by a grant from Bayer AG to the Broad Institute focused on machine learning for clinical trial design. Dr. Ellinor is supported by a grant from Bayer AG to the Broad Institute focused on the genetics and therapeutics of cardiovascular diseases. Dr. Ellinor has also served on advisory boards or consulted for Bayer AG, Quest Diagnostics, MyoKardia and Novartis. Remaining authors report no disclosures.

