## Supplementary Notes and Figures for "Genetic Analysis of Right Heart Structure and Function in 40,000 People"

### Supplementary Material

**Supplementary Note** 1
Deep learning quality assessment 1
**Supplementary Figures** 3
Supplementary Figure 1: Sample flow diagram 3
Supplementary Figure 2: Right heart trait distributions 4
Supplementary Figure 3: Cardiovascular magnetic resonance imaging trait correlation matrix 5
Supplementary Figure 4: Genetic correlation between cardiovascular imaging phenotypes 6
Supplementary Figure 5: Manhattan plots of all 16 traits 7
Supplementary Figure 6: Correlation of lead SNPs across cardiovascular phenotypes 8
A: 8
B: 9
Supplementary Figure 7: BAG3 locus for RVESV and LVESV 11
Supplementary Figure 8: TTN locus for RVESV and LVESV 12
Supplementary Figure 9: GATA4 locus for RVESV and LVESV 13
Supplementary Figure 10: OBSCN locus for RVESV and LVESV 14
Supplementary Figure 11: Chamber-specific cell type enrichment 15
Supplementary Figure 12: Pulmonary artery diameter among carriers and noncarriers of AAGAB loss-of-function variants 16
Supplementary Figure 13: Gene sets enriched near GWAS loci 17
Supplementary Figure 14: JUP locus for RVESV and LVESV 18

### Supplementary Note

#### Deep learning quality assessment

The four-chamber long axis deep learning model output was compared to a held-out test set of 40 additional manually annotated images (by JPP) that were not used for training or validation. The Dice coefficient of the deep learning model output was 0.94 ± 0.03 (representing mean ± standard deviation [SD]) for the right atrial blood pool and 0.94 ± 0.02 for the right ventricular blood pool. The short axis deep learning model output was assessed against 57 additional short axis test set images (manually annotated by JPP). The average Dice coefficient of the deep learning model output was 0.89 ± 0.21 for the right ventricular blood pool and 0.96 ± 0.16 for the pulmonary artery blood pool.

From those values, we observed that, despite similar point estimates, the short axis measurements had greater variance in the Dice score compared to the four-chamber long axis measurements, despite the short axis model having more training data. We took a series of steps to investigate.

A second cardiologist (VN) annotated 300 randomly selected short axis images. The first annotator (JPP) annotated 50 additional images from the set of 300. The Dice scores for the RV annotations comparing the annotators with one another and with the deep learning models are shown in **Supplementary Table 11**. Across all comparisons, we observed the same pattern of high Dice score standard deviation across images for the right ventricle.

Finally, we looked at properties of all available images on which the models were successfully applied (24,378,930 short axis images and 2,296,549 four-chamber long axis images). On average, the short axis images contained 39,600 ± 4,500 pixels, whereas the four-chamber long axis images contained 35,200 ± 3,700 pixels. This yielded a larger coefficient of variation (standard deviation divided by mean) for the number of pixels in short axis images than in four-chamber long axis images (0.114 vs 0.105). In addition, the portion of each image occupied by the structures of interest was smaller in the short axis images than in the four-chamber long axis images. The right ventricular blood pool was measured in both views and offers a direct comparison: 0.68±0.79% of pixels in the short axis and 1.78±0.53% of pixels in the four-chamber long axis view were annotated as belonging to the right ventricle.

We concluded that both (a) the portion of the image occupied by structures of interest (0.68% for short axis vs 1.78% for four-chamber long axis), and (b) the variability of how much of the image is occupied by those structures (coefficient of variation 1.16 for short axis and 0.30 for four-chamber long axis) likely contribute to the greater variation in Dice score for the annotated structures of the short axis images. Of note, when the Dice scores were weighted by the total number of pixels attributed to the RV (represented by the “Weighted Dice” columns in **Supplementary Table 11**), we found that the variance was lower, consistent with greater agreement in the images that ultimately contribute more pixels to the RV estimates in the downstream Poisson reconstruction models.

### Supplementary Figures

#### Supplementary Figure 1: Sample flow diagram

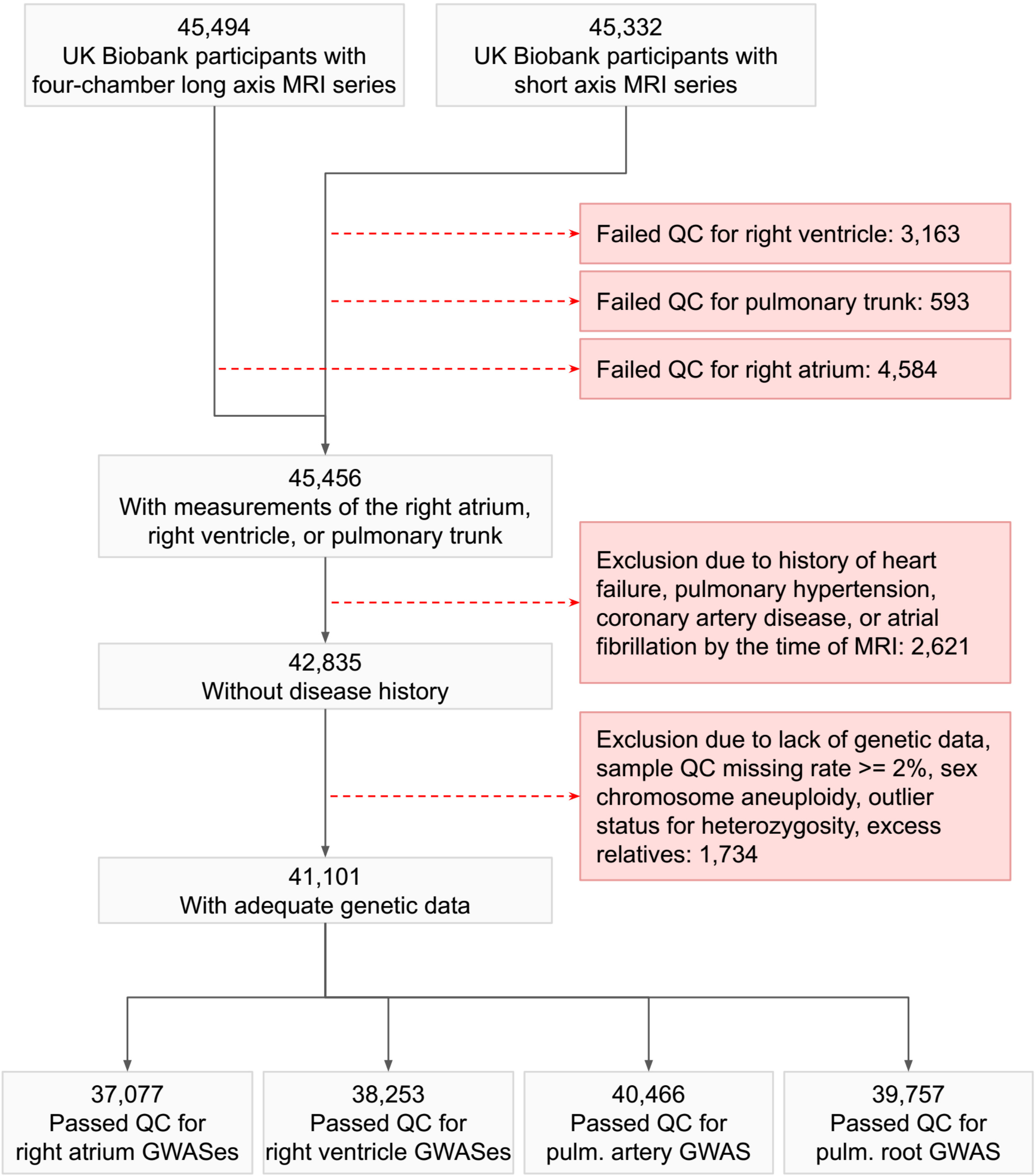

### Supplementary Figure 2: Right heart trait distributions

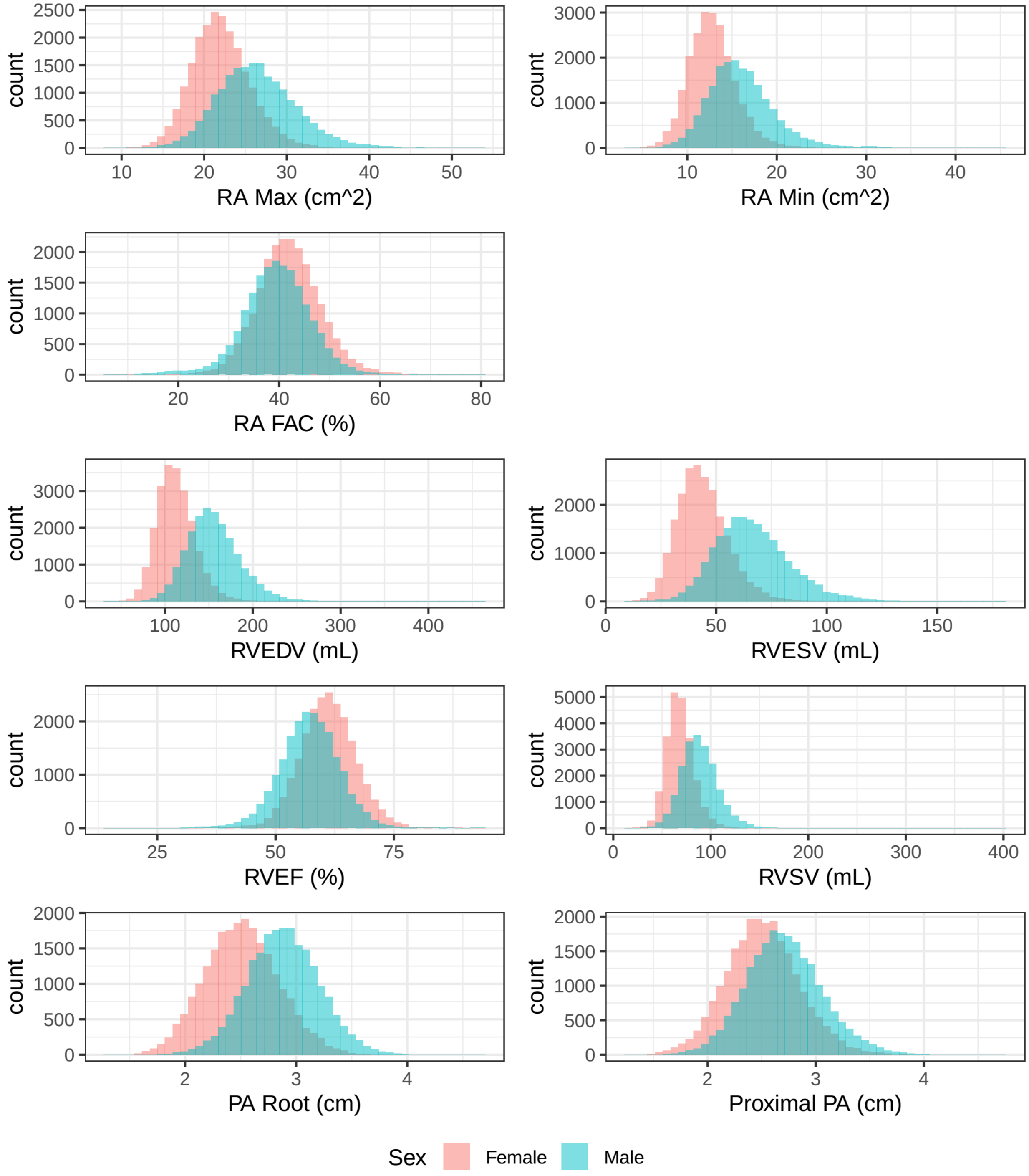


Histograms are shown for each trait in its natural units (**x axis**) with the number of individuals in each bucket on the **y axis**. Measurements for women are depicted in red and for men in turquoise.

### Supplementary Figure 3: Cardiovascular magnetic resonance imaging trait correlation matrix

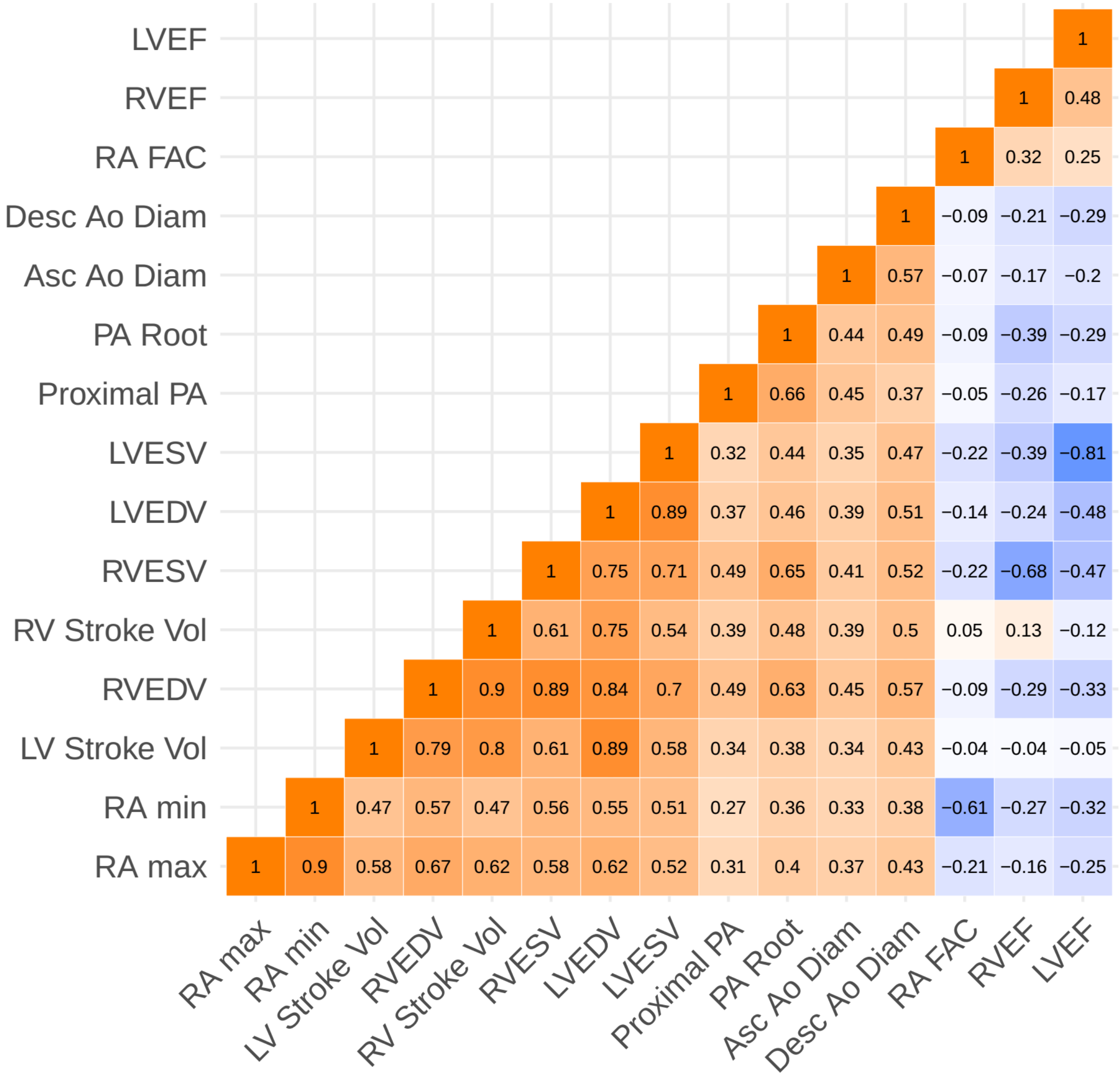


The same traits are shown on the x and y axes. The trait intersections represent the correlation (r) between the traits.

#### Supplementary Figure 4: Genetic correlation between cardiovascular imaging phenotypes

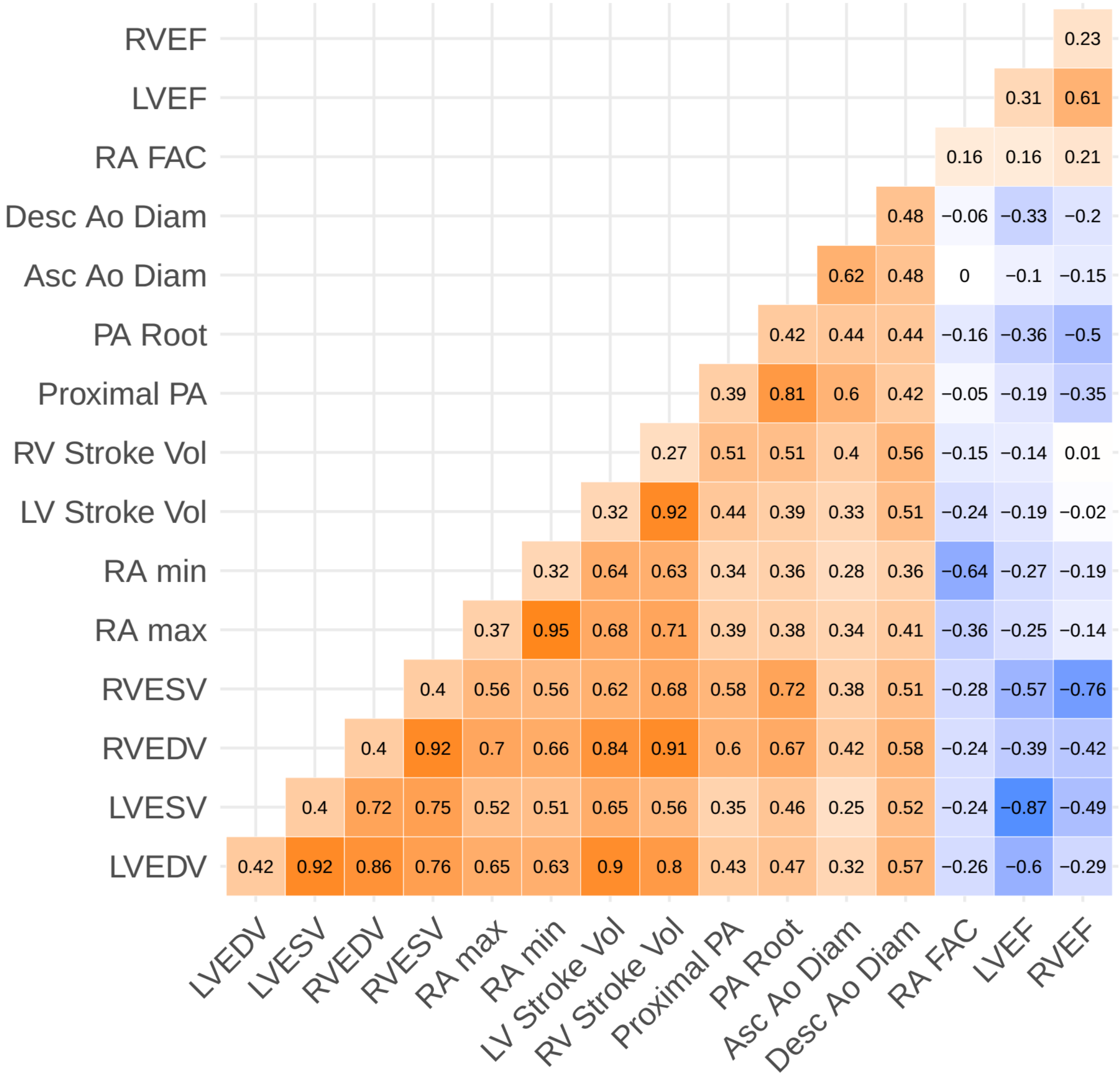


The same traits are shown on the x and y axes. The trait intersections represent the genetic correlation (rg) between the traits. Each trait's heritability (h2g) is shown along the diagonal.

### Supplementary Figure 5: Manhattan plots of all 16 traits

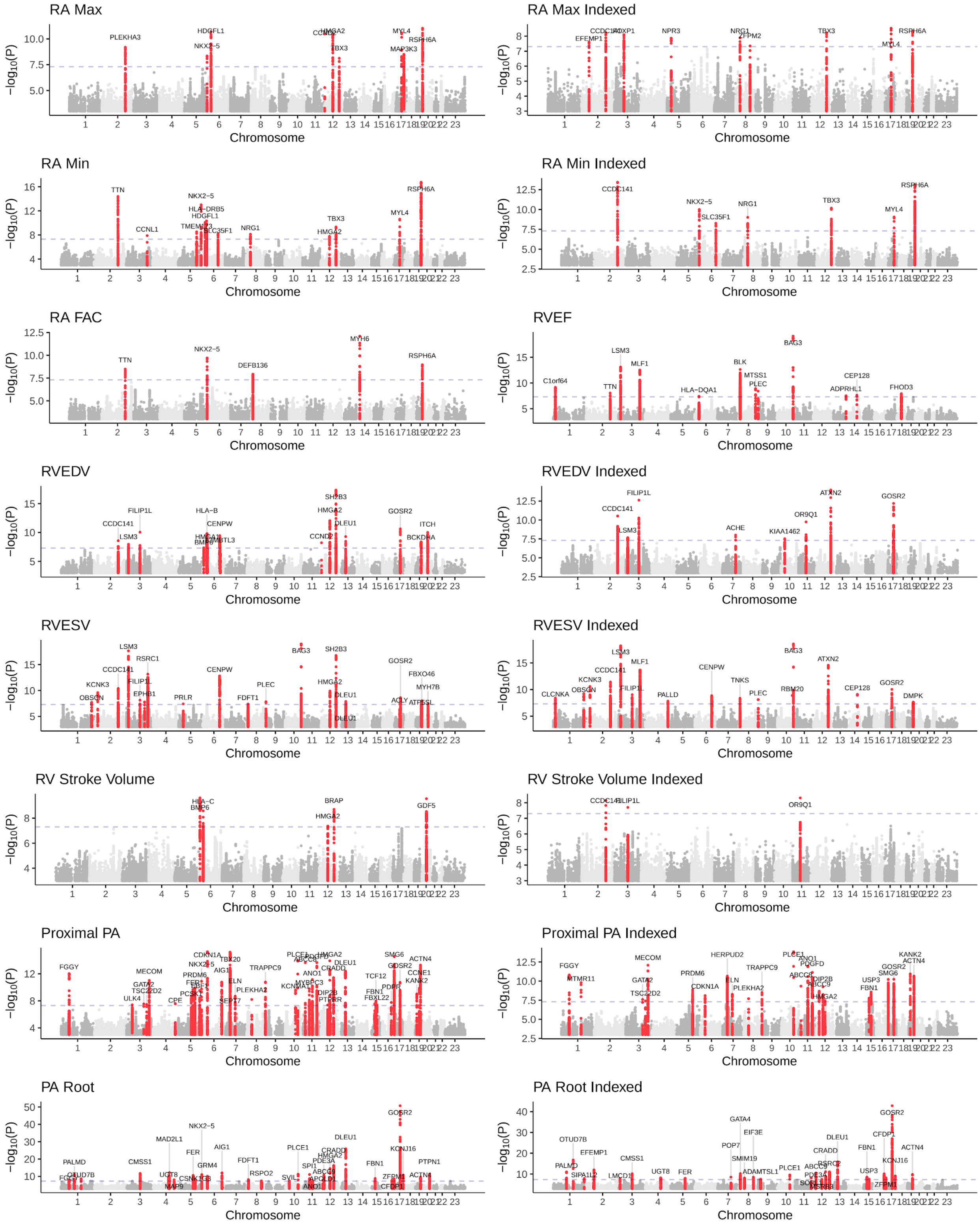


Manhattan plots are depicted for each of the 16 phenotypes (including 9 raw phenotypes and 7 body surface area-indexed phenotypes). The **x axes** represent the chromosome and position.

The **y axes** represent the -$\log_{10}$(BOLT LMM P). Loci that contain SNPs with P < 5E-08 are colored red and labeled with the name of the nearest gene.

#### Supplementary Figure 6: Correlation of lead SNPs across cardiovascular phenotypes

A:

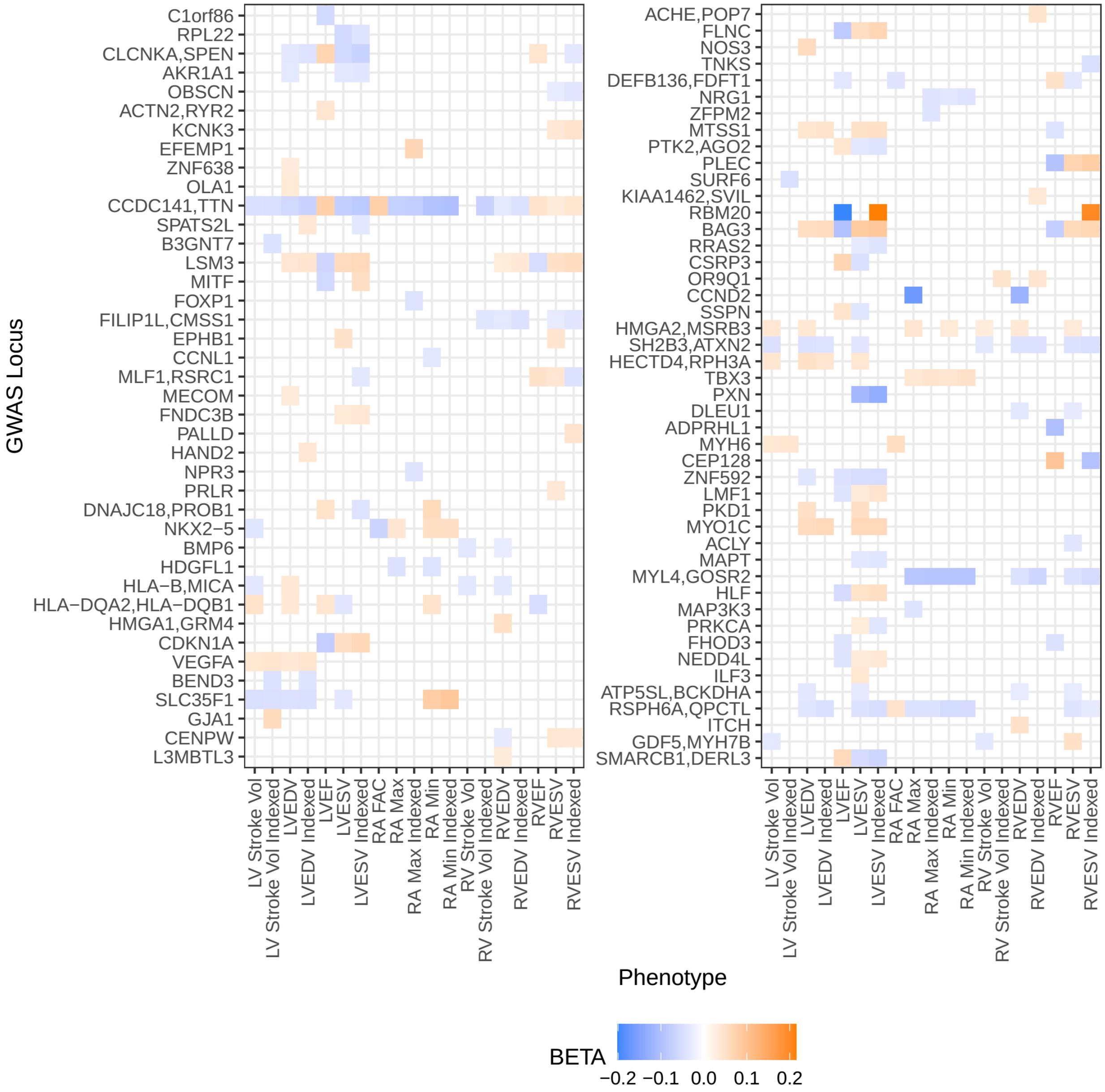

B:

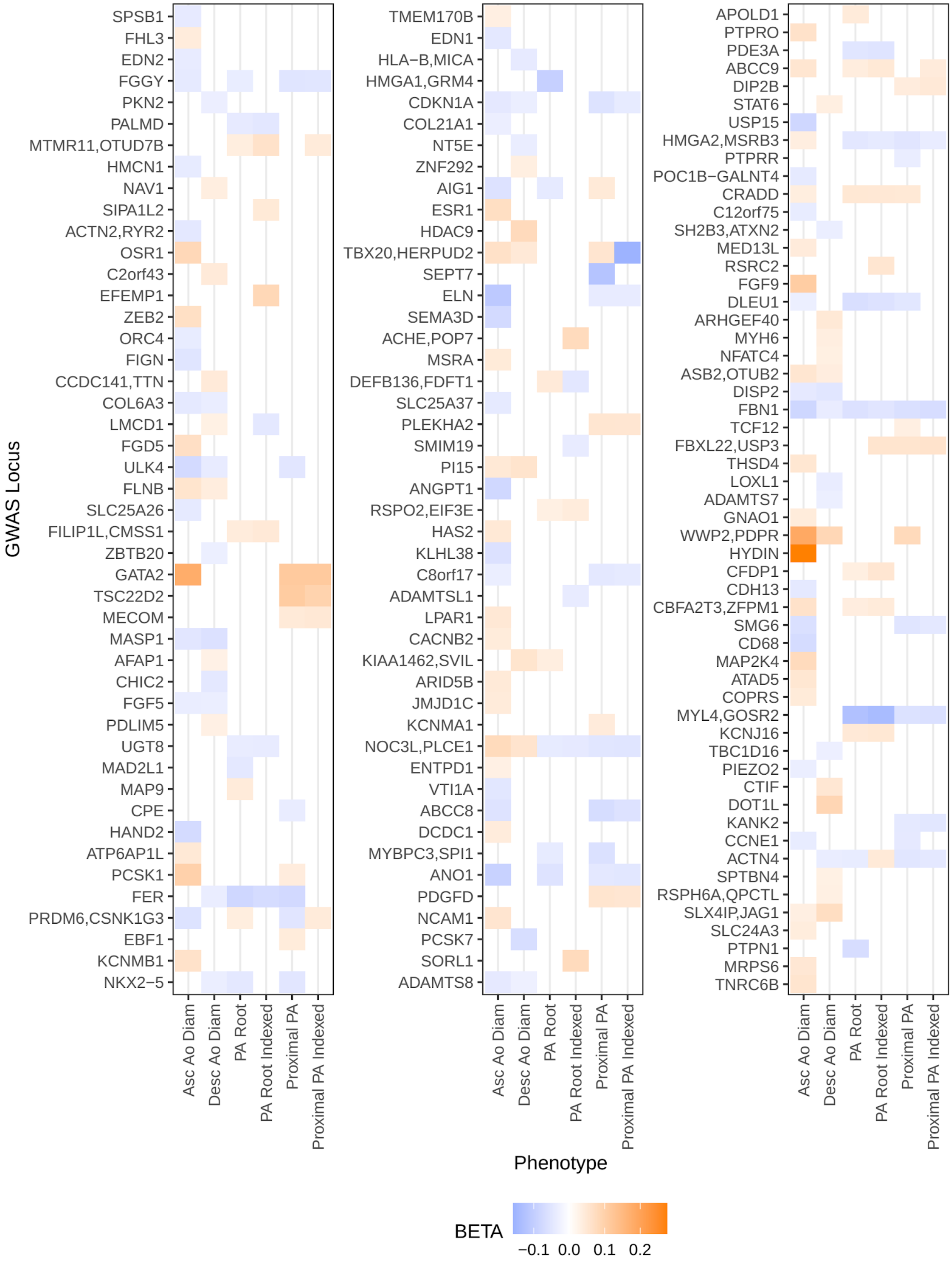
GWAS Locus
SPSB1
FHL3
EDN2
FGGY
PKN2
PALMD
MTMR11,OTUD7B
HMCN1
NAV1
SIPA1L2
ACTN2,RYR2
OSR1
C2orf43
EFEMP1
ZEB2
ORC4
FIGN
CCDC141,TTN
COL6A3
LMCD1
FGD5
ULK4
FLNB
SLC25A26
FILIP1L,CMSS1
ZBTB20
GATA2
TSC22D2
MECOM
MASP1
AFAP1
CHIC2
FGF5
PDLIM5
UGT8
MAD2L1
MAP9
CPE
HAND2
ATP6AP1L
PCSK1
FER
PRDM6,CSNK1G3
EBF1
KCNMB1
NKX2−5
TMEM170B
EDN1
HLA−B,MICA
HMGA1,GRM4
CDKN1A
COL21A1
NT5E
ZNF292
AIG1
ESR1
HDAC9
TBX20,HERPUD2
SEPT7
ELN
SEMA3D
ACHE,POP7
MSRA
DEFB136,FDFT1
SLC25A37
PLEKHA2
SMIM19
PI15
ANGPT1
RSPO2,EIF3E
HAS2
KLHL38
C8orf17
ADAMTSL1
LPAR1
CACNB2
KIAA1462,SVIL
ARID5B
JMJD1C
KCNMA1
NOC3L,PLCE1
ENTPD1
VTI1A
ABCC8
DCDC1
MYBPC3,SPI1
ANO1
PDGFD
NCAM1
PCSK7
SORL1
ADAMTS8
APOLD1
PTPRO
PDE3A
ABCC9
DIP2B
STAT6
USP15
HMGA2,MSRB3
PTPRR
POC1B−GALNT4
CRADD
C12orf75
SH2B3,ATXN2
MED13L
RSRC2
FGF9
DLEU1
ARHGEF40
MYH6
NFATC4
ASB2,OTUB2
DISP2
FBN1
TCF12
FBXL22,USP3
THSD4
LOXL1
ADAMTS7
GNAO1
WWP2,PDPR
HYDIN
CFDP1
CDH13
CBFA2T3,ZFPM1
SMG6
CD68
MAP2K4
ATAD5
COPRS
MYL4,GOSR2
KCNJ16
TBC1D16
PIEZO2
CTIF
DOT1L
KANK2
CCNE1
ACTN4
SPTBN4
RSPH6A,QPCTL
SLX4IP,JAG1
SLC24A3
PTPN1
MRPS6
TNRC6B
Asc Ao Diam
Desc Ao Diam
PA Root
PA Root Indexed
Proximal PA
Proximal PA Indexed
Phenotype
BETA
−0.1 0.0 0.1 0.2

Cardiac (**Panel A**) and great vessel (**Panel B**) loci are plotted. Each locus is identified by the name of the nearest gene to one or more SNPs. Effect size of the rank-based inverse normalized phenotypes at the locus (**y-axis**) for a phenotype (**x-axis**) are identified by color; where no genome-wide significant association for a trait at a locus, no color is plotted. Effect directions are matched based on minor allele frequency. Note that, as described in the main text, the lead SNPs in some cases are uncorrelated between phenotypes, in which case the effect direction is not meaningful.

### Supplementary Figure 7: *BAG3* locus for RVESV and LVESV

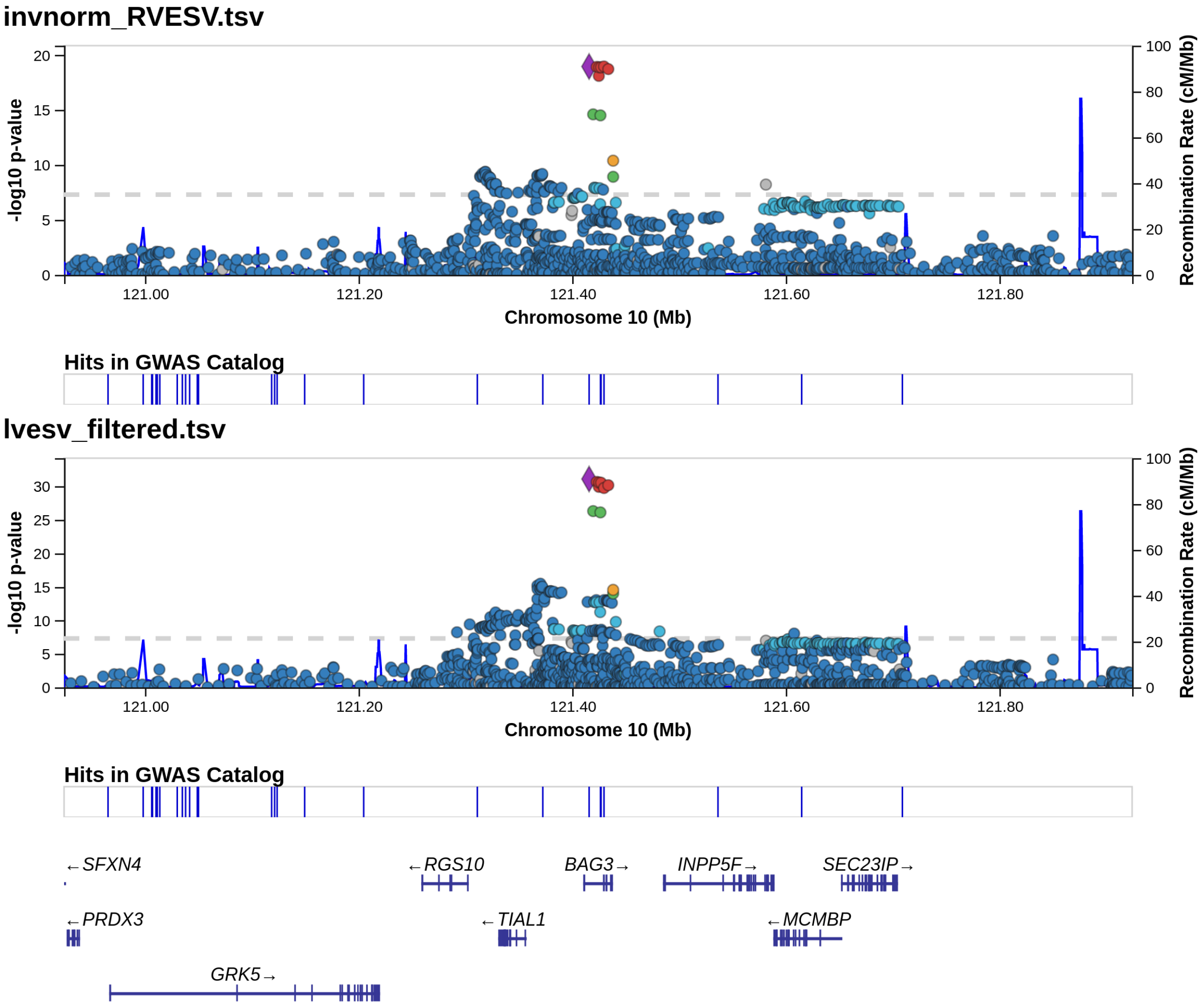

### Supplementary Figure 8: *TTN* locus for RVESV and LVESV

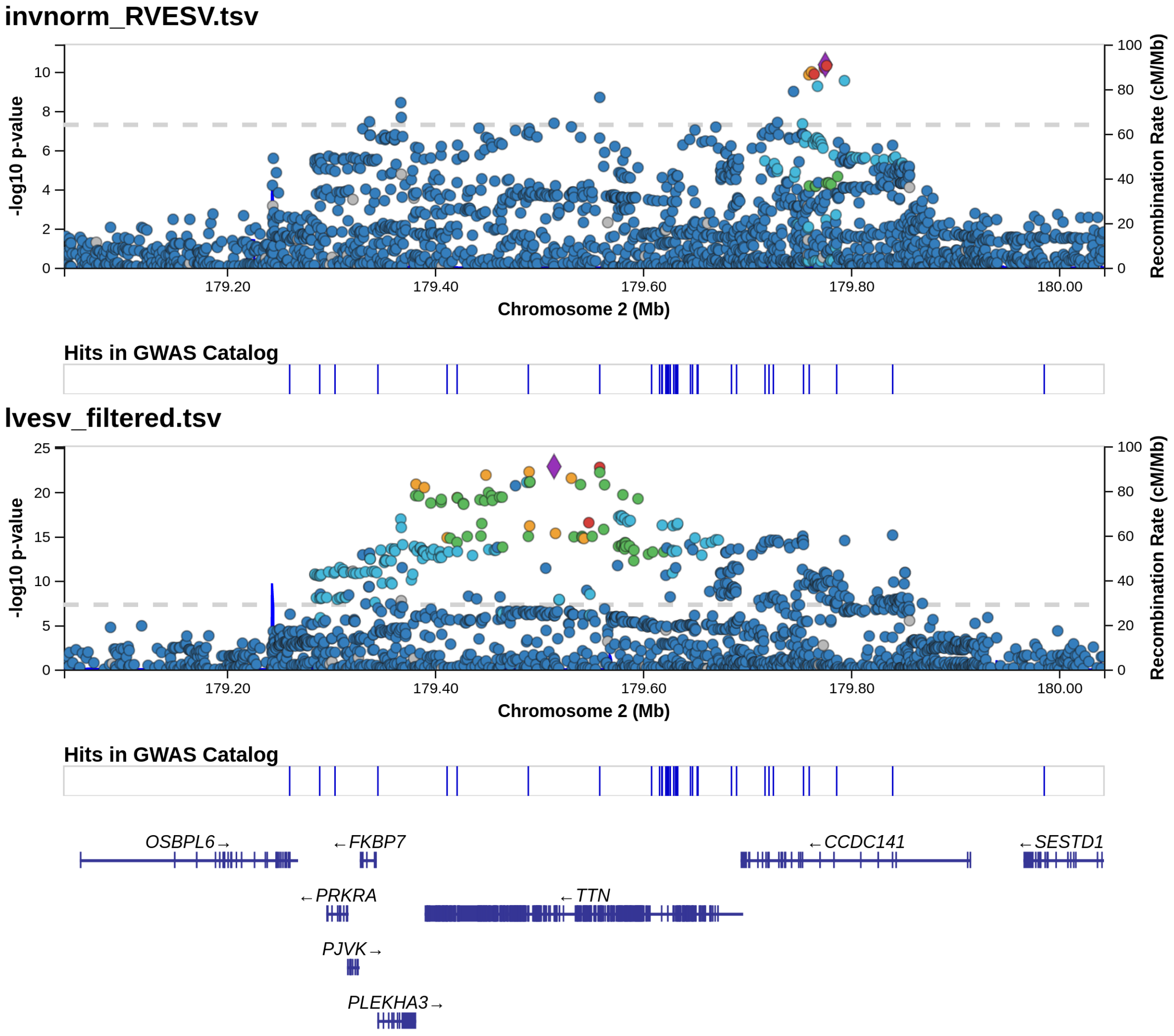

### Supplementary Figure 9: *GATA4* locus for RVESV and LVESV

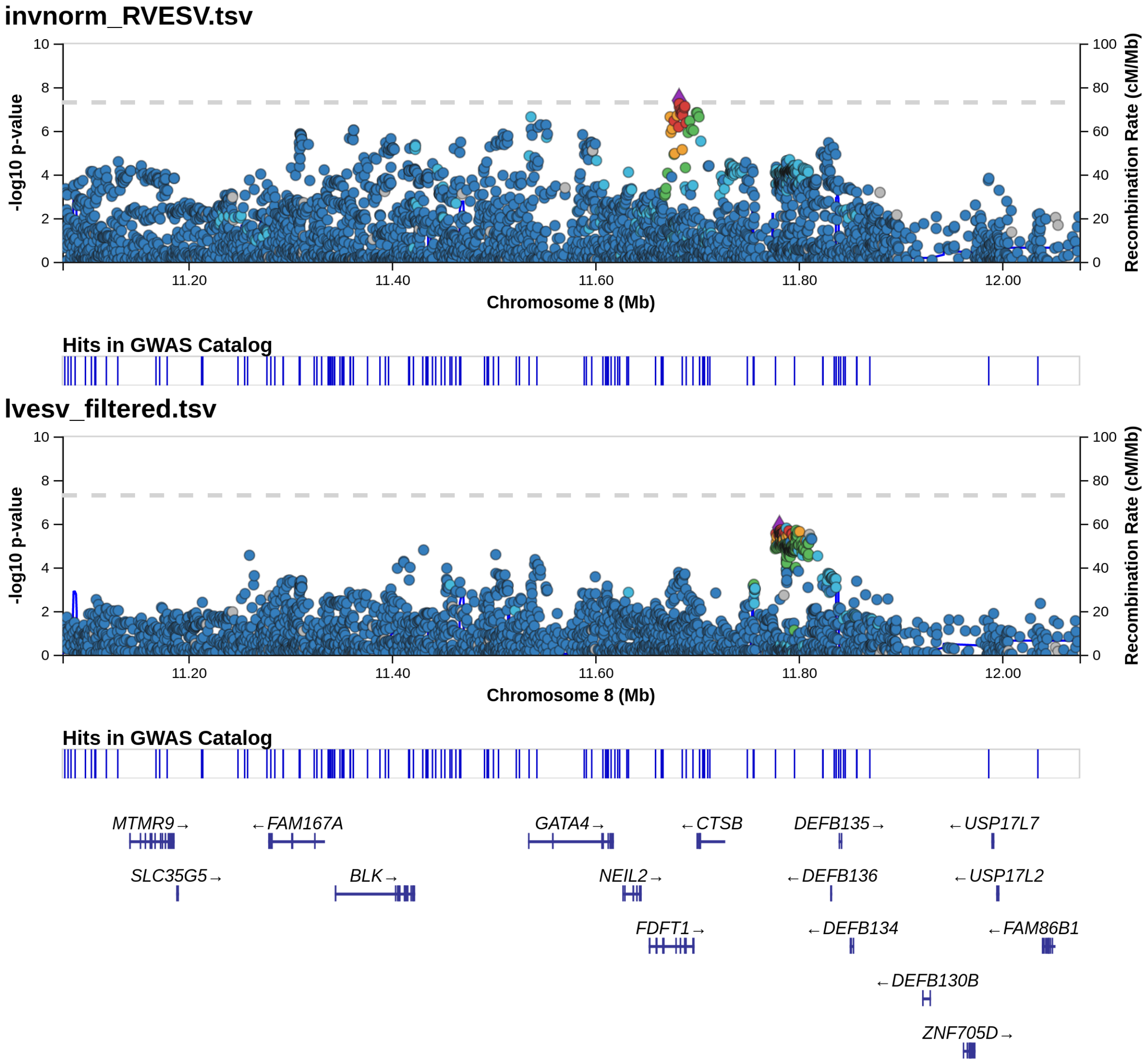

### Supplementary Figure 10: *OBSCN* locus for RVESV and LVESV

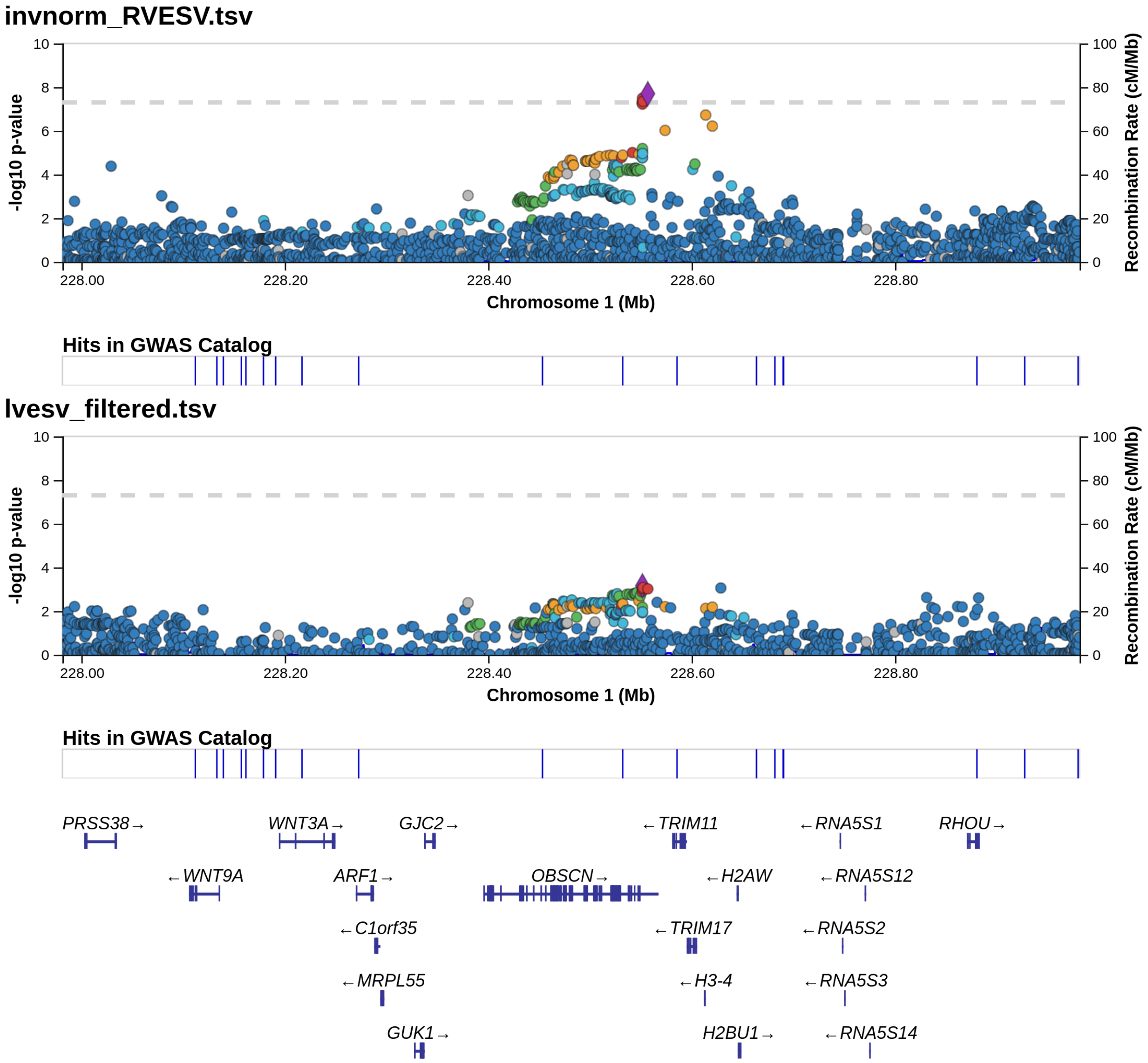

#### Supplementary Figure 11: Chamber-specific cell type enrichment

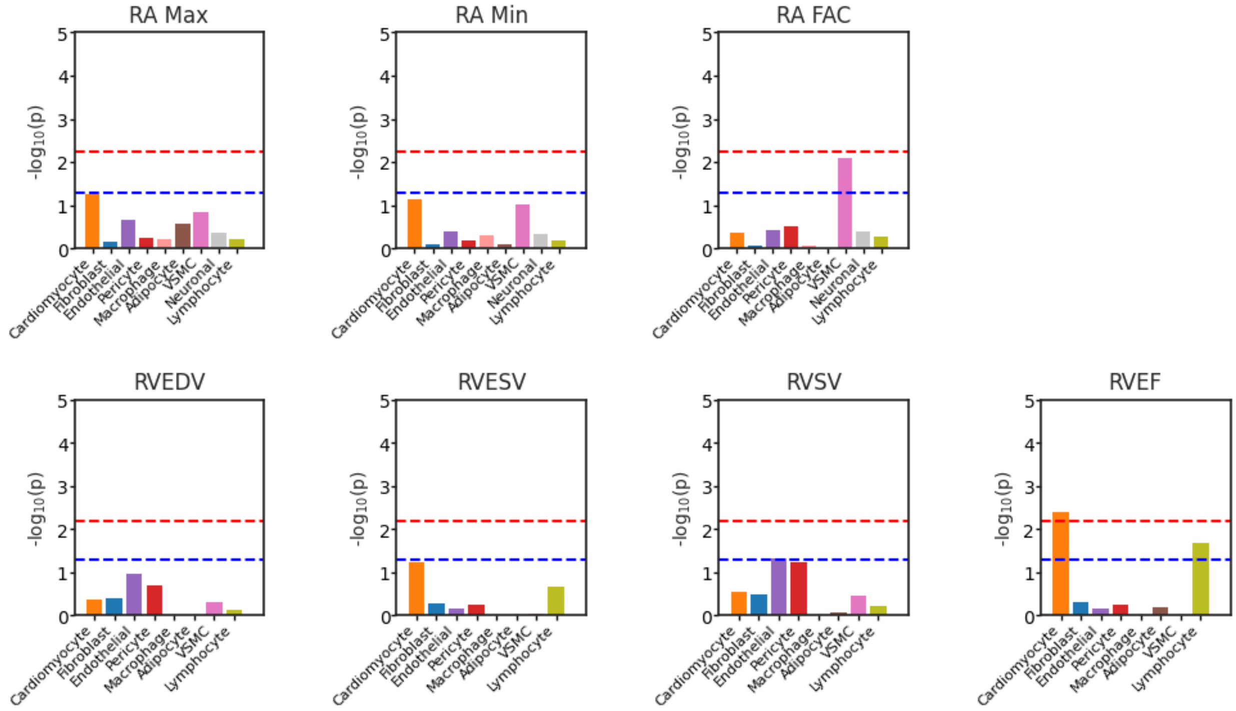


For the three primary right atrial phenotypes and four primary right ventricular phenotypes, stratified LD-score regression was performed. For the right atrial phenotypes, right atrial expression data was used, and for the right ventricular phenotypes, right ventricular expression data was used. The blue dashed line represents nominal significance while the red dashed line represents trait-wise significance.

Supplementary Figure 12: Pulmonary artery diameter among carriers and noncarriers of *AAGAB* loss-of-function variants

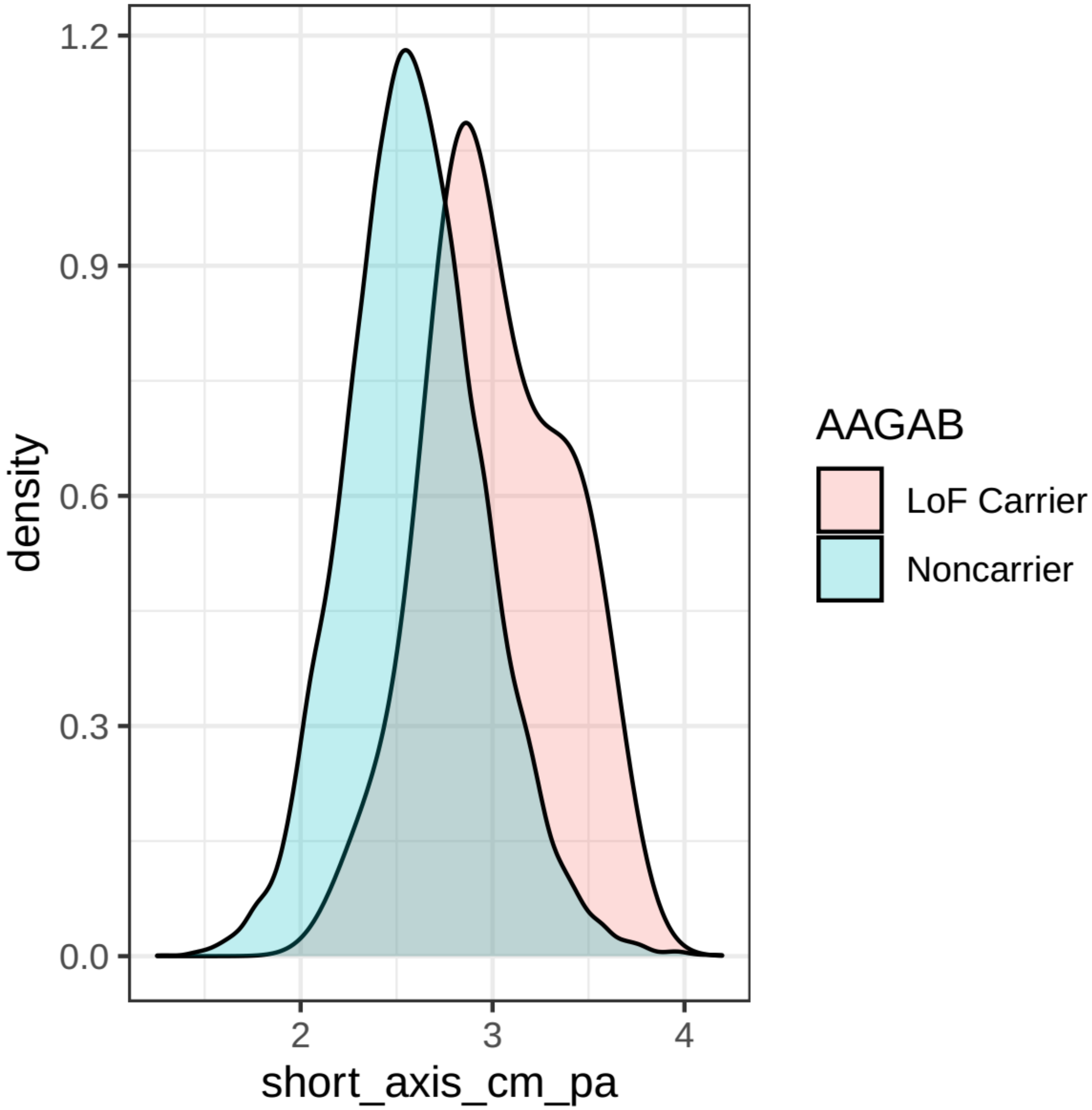

#### Supplementary Figure 13: Gene sets enriched near GWAS loci

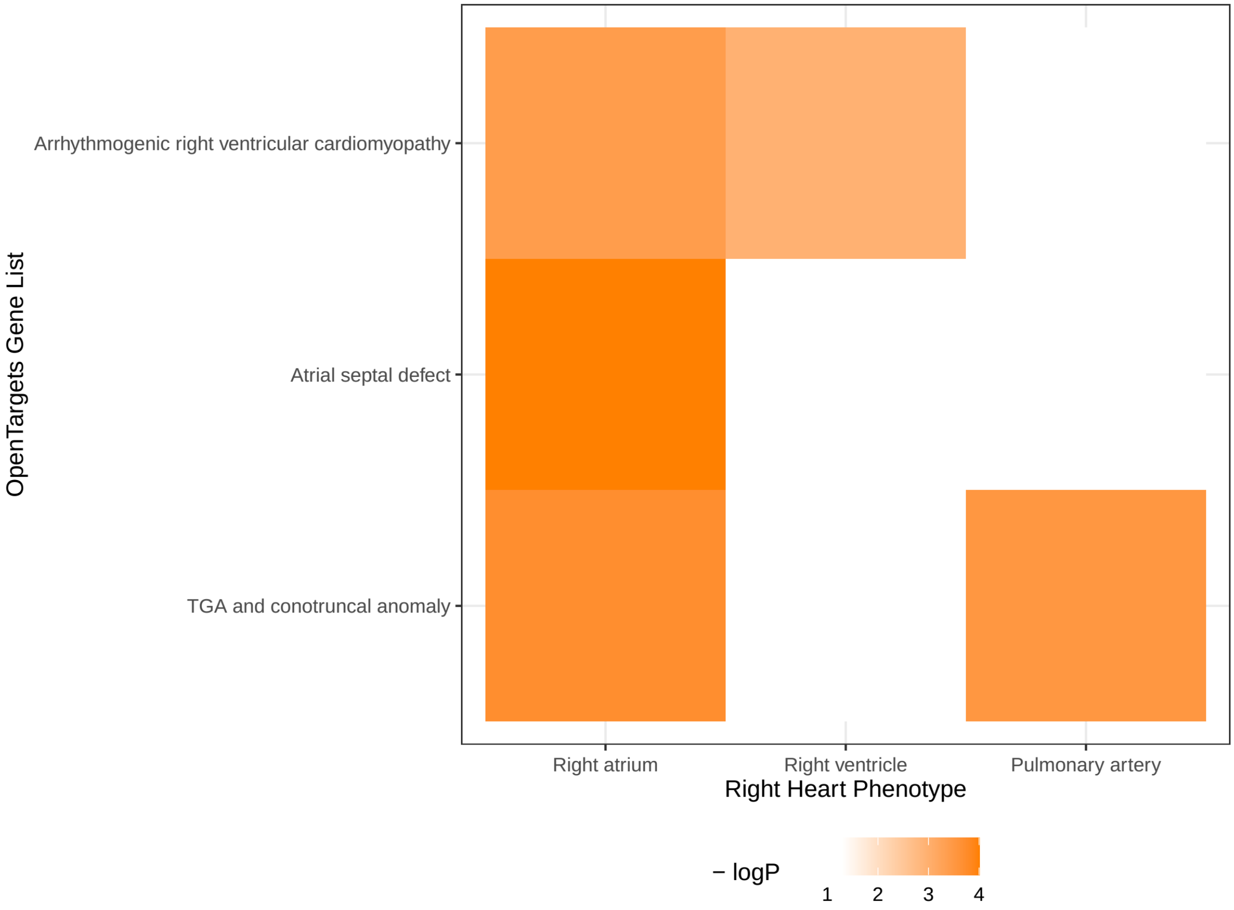

Supplementary Figure 14: *JUP* locus for RVESV and LVESV

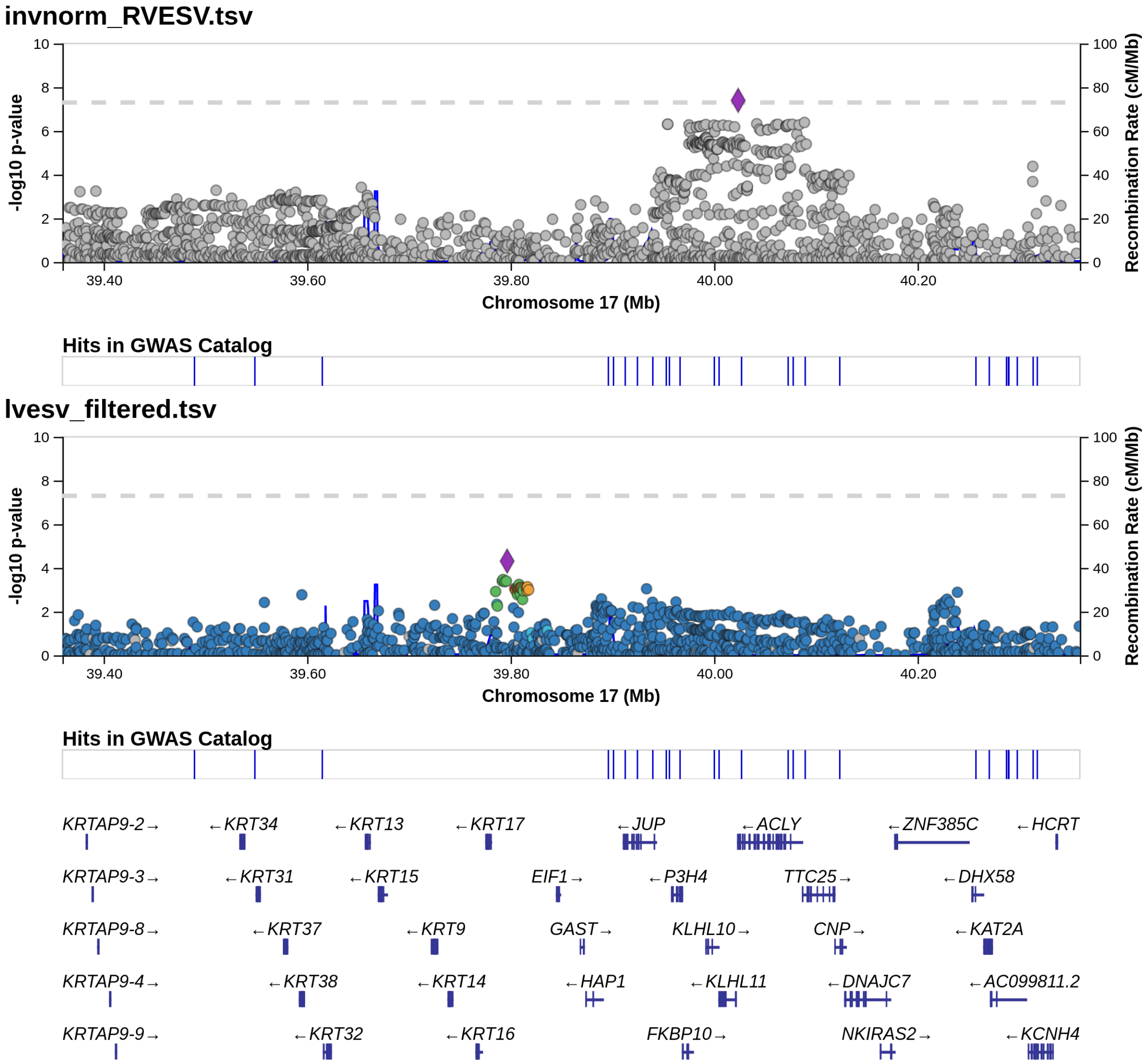
